# Normative and mechanistic model of an adaptive circuit for efficient encoding and feature extraction

**DOI:** 10.1101/2021.09.24.461723

**Authors:** Nikolai M. Chapochnikov, Cengiz Pehlevan, Dmitri B. Chklovskii

**Affiliations:** Flatiron Institute, Simons Foundation, New York, NY, USA; Department of Neurology, New York University School of Medicine, New York, NY, USA; John A. Paulson School of Engineering and Applied Sciences, Harvard University, Cambridge, MA, USA; Neuroscience Institute, New York University School of Medicine, New York, NY, USA

## Abstract

One major question in neuroscience is how to relate connectomes to neural activity, circuit function, and learning. We offer an answer in the peripheral olfactory circuit of the *Drosophila* larva, composed of olfactory receptor neurons (ORNs) connected through feedback loops with interconnected inhibitory local neurons (LNs). We combine structural and activity data and, using a holistic normative framework based on similarity-matching, we propose a biologically plausible mechanistic model of the circuit. Our model predicts the ORN → LN synaptic weights found in the connectome and demonstrate that they reflect correlations in ORN activity patterns. Additionally, our model explains the relation between ORN → LN and LN – LN synaptic weight and the arising of different LN types. This global synaptic organization can autonomously arise through Hebbian plasticity, and thus allows the circuit to adapt to different environments in an unsupervised manner. Functionally, we propose LNs extract redundant input correlations and dampen them in ORNs, thus partially whitening and normalizing the stimulus representations in ORNs. Our work proposes a comprehensive framework to combine structure, activity, function, and learning, and uncovers a general and potent circuit motif that can learn and extract significant input features and render stimulus representations more efficient.

**Significance:** The brain represents information with patterns of neural activity. At the periphery, due to the properties of the external world and of encoding neurons, these patterns contain correlations, which are detrimental for stimulus discrimination. We study the peripheral olfactory neural circuit of the Drosophila larva, that preprocesses neural representations before relaying them to higher brain areas. A comprehensive understanding of this preprocessing is, however, lacking. Here, we propose a mechanistic and normative framework describing the function of the circuit and predict the circuit’s synaptic organization based on the circuit’s input neural activity. We show how the circuit can autonomously adapt to different environments, extracts stimulus features, and decorrelate and normalize input representations, which facilitates odor discrimination downstream.

## Introduction

Thanks to technological advances in connectomics (Eichler et al., 2017; Scheffer et al., 2020) and neural population activity imaging (Aimon et al., 2019), more and more neural circuits will soon be characterized anatomically and physiologically at unprecedented scale and detail. However, it is not clear what insights can be obtained from combining such datasets and how to use them to advance our understanding of brain computation. To address this, we focus on the peripheral olfactory system of the *Drosophila* larva - a small and genetically tractable circuit for which a connectivity (Berck et al., 2016) and comprehensive activity imaging (Si et al., 2019) datasets are already available.

This circuit is an analogous, but simpler version of the well-studied olfactory circuit in adult flies and vertebrates (Wilson, 2013). It contains 21 olfactory receptor neurons (ORNs), each expressing a different receptor type with a different odor sensitivity profile (**Fig. 1A**). ORN axons are reciprocally connected to a web of multiple interconnected inhibitory local neurons (LNs) through feedforward excitation and feedback inhibition. The connectome dataset contains not just the presence or absence of a connection between two neurons but also the number of synaptic contacts in parallel (Berck et al., 2016), which is an estimate of the connection strength, since synaptic contacts do not vary significantly in size in the *Drosophila* (Scheffer et al., 2020). The activity dataset contains the responses of ORNs to 34 odors at 5 dilutions (**Fig. 2A**) and has been obtained by imaging Ca^2+^ concentration in their somas (Si et al., 2019).

**Fig. 1.**
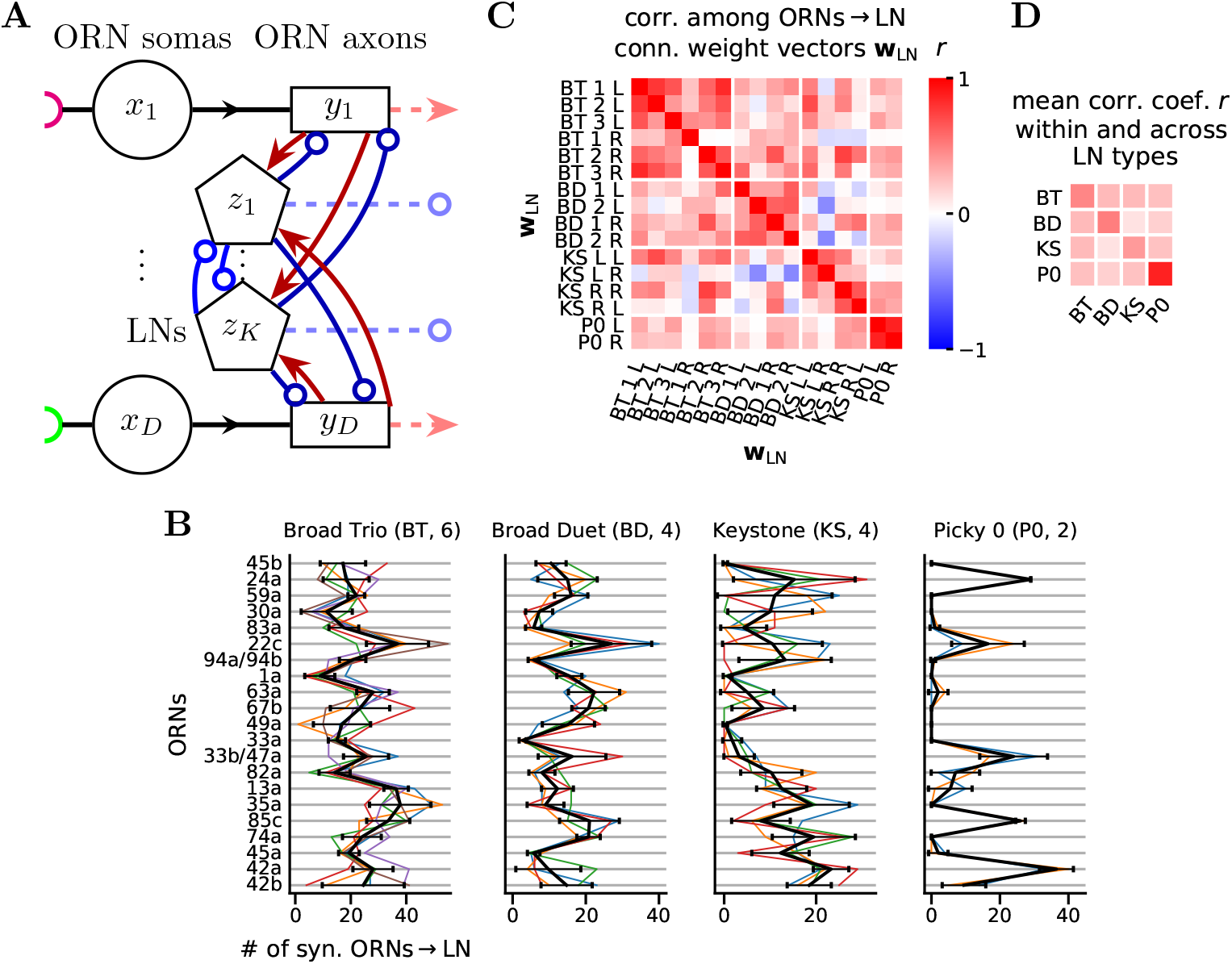
Circuit connectivity and LN types. **A** Scheme of the ORN-LN circuit. Each of the *D* ORNs is depicted as a two-compartment unit with a soma (circle) and an axonal terminal (rectangle). The differently colored half circles on the left represent different chemical receptor types. *K* inhibitory local neurons (LNs, pentagons) reciprocally connect with ORN axons and between themselves. ORN axons and LNs transmit information further downstream (dashed lines). Red lines with arrowheads and blue lines with open circles represent excitatory and inhibitory connections, respectively, *x_i_, y_i_*, and *z_i_* represent the activity of ORN somas, axons, and LNs, respectively. **B** Feedforward ORNs → LN connection weight vectors, **w**_LN_ (colored lines), and average feedforward ORNs → LN type connection weight vectors, **w**_LNtype_ (black lines, mean ± s.d.) for each LN type (see also **Fig. S2A**). **C** Correlation coefficients *r* between all **w**_LN_. L: left, R: right. KS L R is the Keystone with the soma positioned on the left side of the larva, connecting with the ORNs of the right side, and vice-versa for KS R L. Since Picky 0 receives synaptic input mainly on the dendrite, here we only use the connections synapsing onto the dendrite. **D** Average rectified correlation coefficient 〈*r*_+_〉 (*r*_+_:= max[0, *r*]) between LN types calculated by averaging the rectified values from (**C**) in each rectangle with white border, excluding the diagonal entries of the full matrix.

**Fig. 2.**
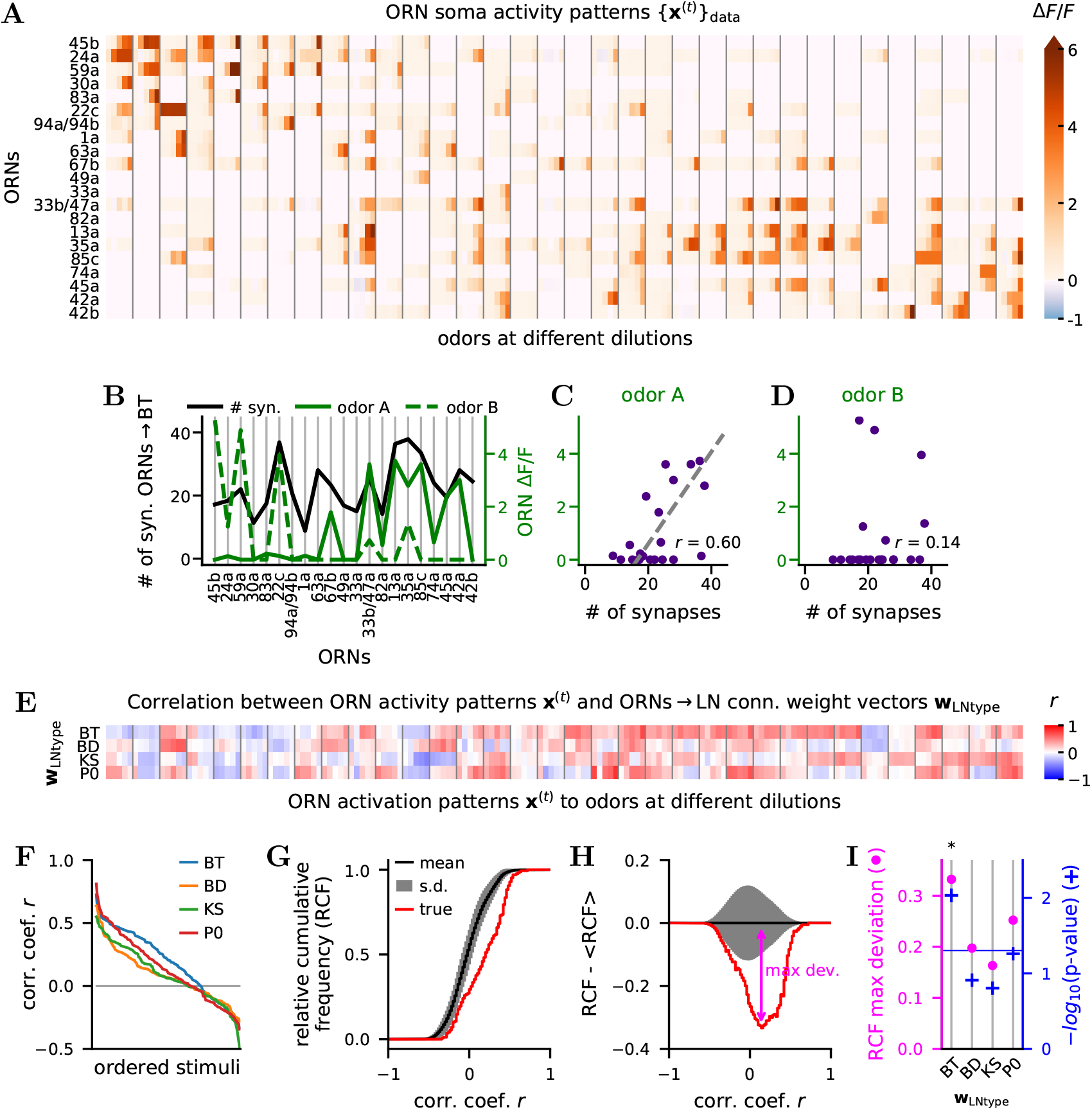
Alignment of ORNs → LN connectivity weight vectors with odor representations in ORN activity. **A** Activity patterns {**x**^(*t*)^}_data_ at ORN soma in response to 34 odors at 5 dilutions from Si et al., 2019. Different odors are separated by vertical gray lines. For each odor, there are 5 columns corresponding to 5 dilutions: 10^−8^,…, 10^−4^. See **Fig. S3** for odor labels and scaled **x**^(*t*)^. **B w**_BT_ superimposed with ORNs activity patterns **x**^(A)^ and **x**^(B)^ in response to the ligands 2-heptanone (odor A) and 2-acetylpyridine (odor B) at dilution 10^−4^. **C-D** Scatter plot representation of (**B**). **w**_BT_ is more strongly tuned to **x**^(A)^ (*r* = 0.6) than to **x**^(B)^ (*r* = 0.14). **E** Correlation coefficients between **w**_LNtype_ with the **x**^(*t*)^ from (**A**) (see also **Fig. S4A**). **F** LN “connectivity tuning curves”: correlation coefficients sorted in decreasing order from (**E**) for each **w**_LNtype_. **G** Red line: relative cumulative frequency (RCF) of the correlation coefficients *r* of the first row of (**E**). Black line and gray band: mean ± s.d. from the RCFs generated by 10,000 instances of shuffling the entries of **w**_BT_. Bin size: 0.02. **H** Same as (**G**) with the mean RCF subtracted. We define the maximum deviation as the maximum negative difference between the true and the mean RCF of correlation coefficients. **I** RCF maximum deviation and log_10_ of false discovery rate (FDR, Benjamini and Hochberg, 1995) adjusted p-values for each **w**_LNtype_ (see also **Fig. S4B**). *: significance with FDR at 5%.

Previous studies addressed the role of the inhibitory feedback provided by LNs in transforming the neural representation of odors from ORN somas to projection neurons (PNs), which are postsynaptic to ORNs. In adult *Drosophila*, this circuit was suggested to perform gain-control and divisive normalization (Olsen et al., 2010; Olsen & Wilson, 2008), which equalizes different odor concentrations and decorrelates input channels. In the zebrafish larva, an analogous circuit was suggested to whiten the input leading to pattern decorrelation which helps odors discrimination downstream (Friedrich, 2013; Wanner & Friedrich, 2020).

However, the underlying mechanistic principles of computation are still not elucidated. For example, whereas different types of LNs have different connectivity patterns with ORNs in the *Drosophila* larva (Berck et al., 2016), the role of different LN types, their multiplicity, and their specific connectivity is not understood. Also, the peripheral olfactory circuit exhibits synaptic plasticity in response to olfactory environment changes (Arenas et al., 2012; Das et al., 2011; Devaud et al., 2001; Sachse et al., 2007; Sudhakaran et al., 2012), but the functional role of such plasticity is unclear.

To address these shortcomings, we use a combination of data analysis and modeling and develop a holistic theoretical framework that links circuit structure, function, activity data, and learning. Our contribution is fourfold. (1) We find that the ORNs → LN synaptic weights vectors reflect features of the independently acquired ORN activity patterns dataset (**Fig. 2, 3, 4**). (2) Building upon the similarity matching framework (Pehlevan et al., 2018), we develop a novel, biologically realistic, normative circuit model incorporating activity-dependent synaptic plasticity. (3) The model, driven by the ORN activity dataset, predicts the following observations in the structural dataset: the ORNs → LN synaptic weights (**Fig. 4**), the emergence of LNs groups (**Fig. 4**), and the relationship between feedforward ORN → LN and lateral LN - LN connection (**Fig. 5**). (4) Using our model, we characterize the circuit computation (**Fig. 6, 7**), and propose that LNs play a dual role in rending the neural representation of odors in ORNs more efficient and extracting useful features that are transmitted downstream. Furthermore, we show that the synaptic weights enabling this computation can be learned by the circuit in an unsupervised manner.

**Fig. 3.**
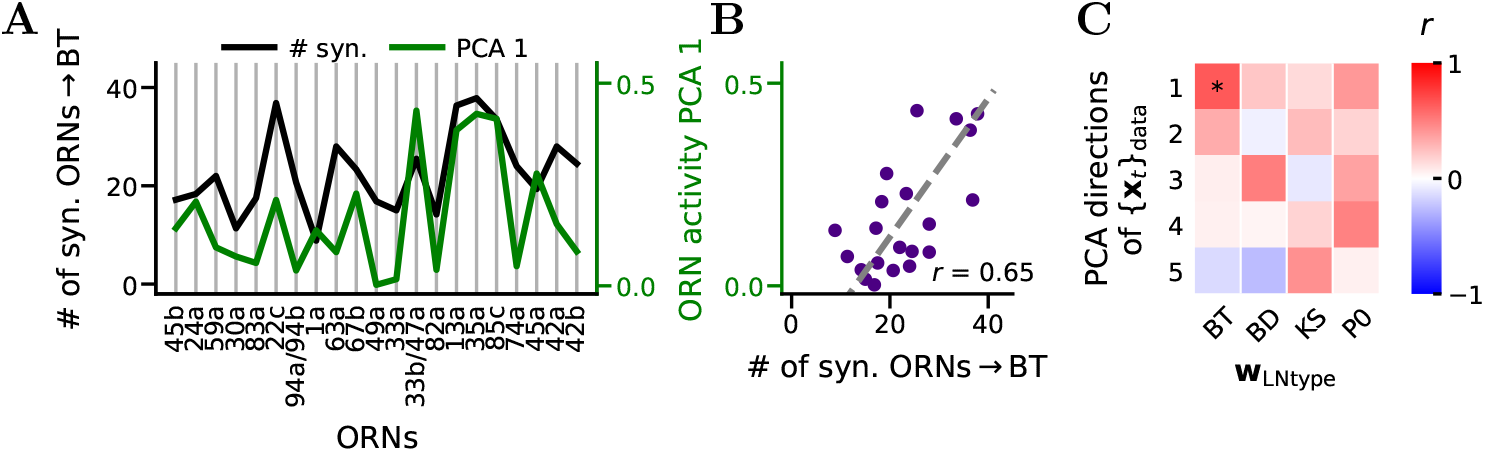
Alignment of w_BT_ with the top PCA direction of ORN activity patterns {**x**^(*t*)^}_data_. **A w**_BT_ superimposed in the 1^st^ PCA direction of {**x**^(*t*)^}_data_ **B** Scatter plot representation of (**A**). **C** Correlation coefficient *r* between the top 5 principal directions of {**x**^(*t*)^}_data_ and the four **w**_LNtype_ (see also **Fig. S6C,D,G**). Two-sided p-values were calculated by shuffling the entries of each **w**_LNtype_. 50,000 permutations used. *: significance with FDR at 5%.

**Fig. 4.**
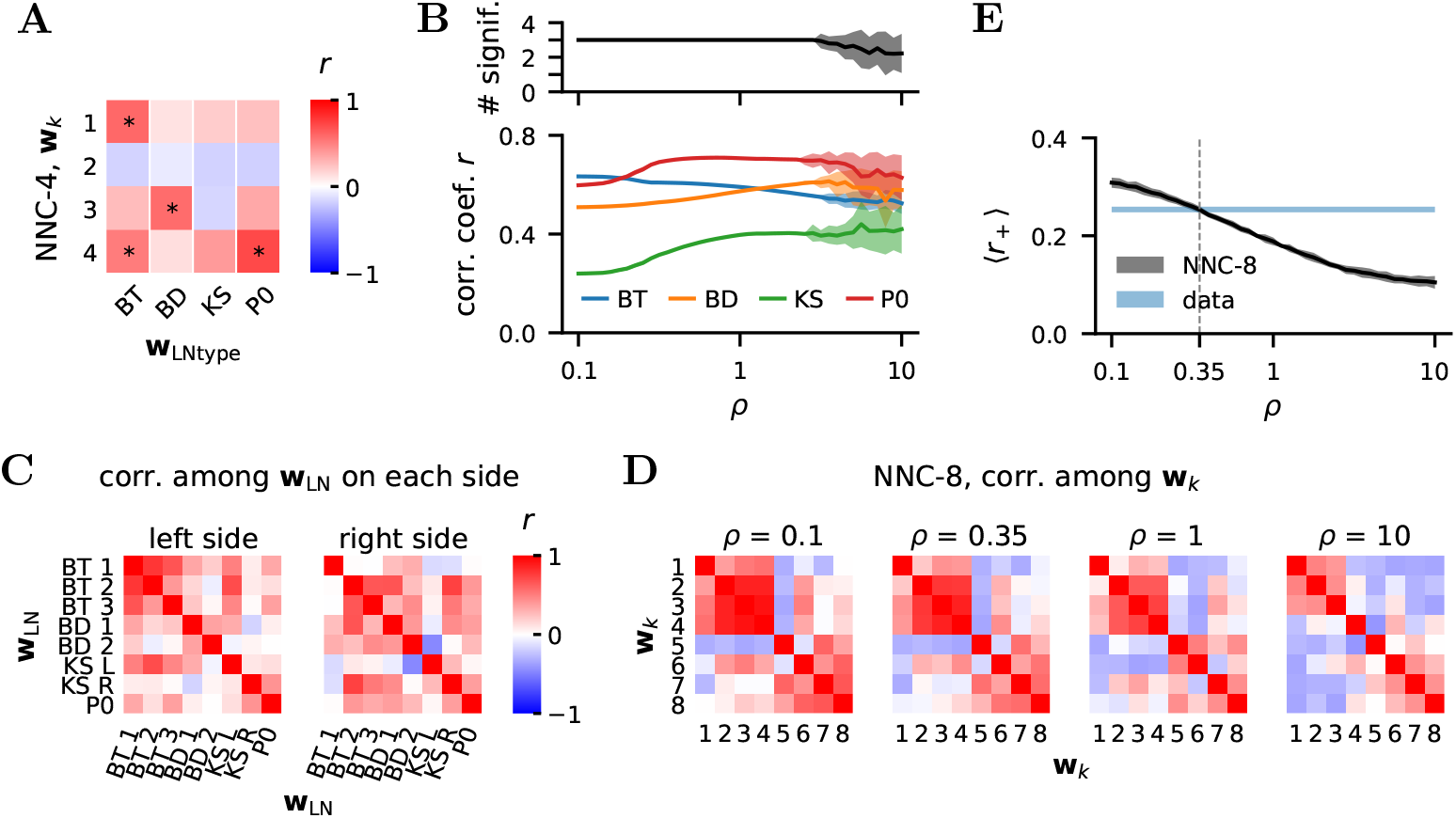
Prediction of the connectivity with the NNC and emergence of LN types. **A** Correlation coefficient *r* between the four **w***_k_* from NNC-4 (*ρ* = 1) and the four **w**_LNtype_ (see also **Fig. S6C,D,F-H**). One-sided p-values were calculated by shuffling the entries of each **w**_LNtype_. 50,000 permutations used. *: significance with FDR at 5%. **B** Bottom: maximum correlation coefficient (mean ± s.d.) of the four **w***_k_* from NNC-4 with the four **w**_LNtype_ for different values of *ρ*. Top: number of **w**_LNtype_ significantly correlated with at last one **w***_k_* from NNC-4 (FDR at 5%). 50 numerical simulations of NNC-4 for each value of *ρ*. **C** Correlation between the **w**_LN_ on the left and right sides of the larva brain. **D** Same as (**C**) for the eight **w***_k_* arising from NNC-8 and with *ρ* = 0.1, 0.35,1,10. **w***_k_* ordered with hierarchical clustering. **E** M ean rectified correlation coefficient 〈*r*_+_〉 from (**C**) (blue band delimited by the value for left and right circuit) and from NNC-8 (black line, mean ± s.d.). One 〈*r*_+_〉 is obtained by averaging all the rectified values in a matrix in (**C**) or (**D**), excluding the diagonal. For the NNC-8 and a given value of *ρ*, we run 50 simulations. Each simulation can give rise to a different set of **w***_k_*, we thus plot the mean ± s.d. of all the 50 〈*r*_+_〉 for a given *ρ*.

**Fig. 5.**
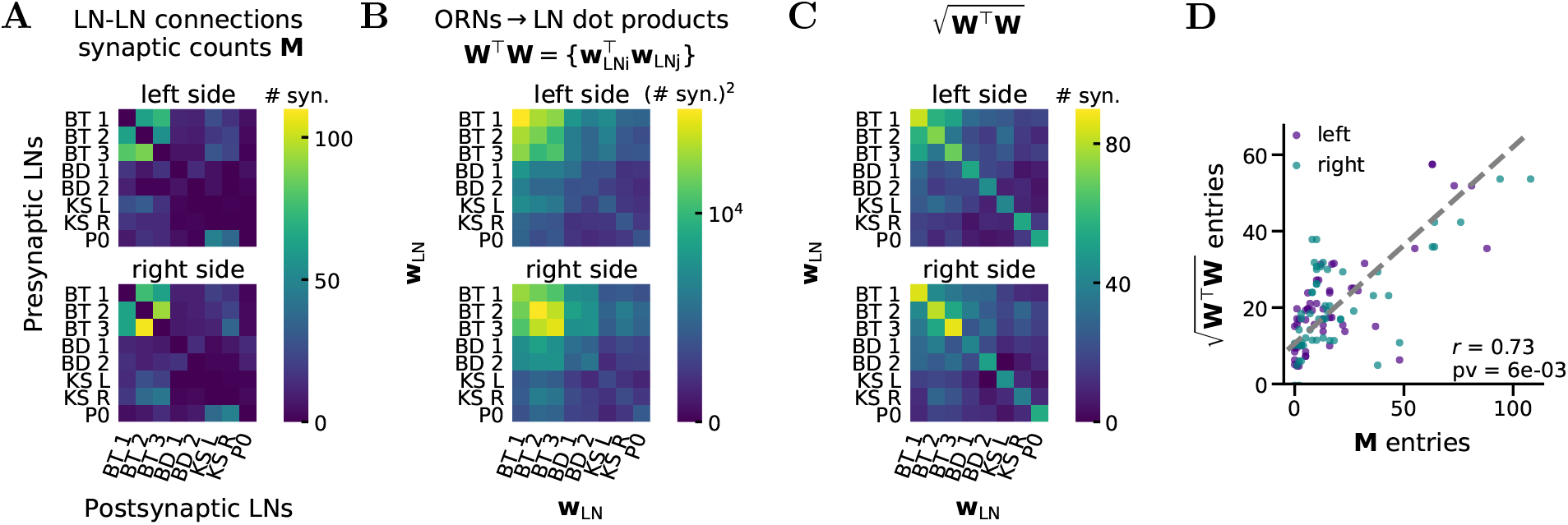
Relation between LN-LN (M) and ORNs → LN (W) synaptic counts in the connectome reconstruction. **A** LN-LN connections synaptic counts **M** on the left and right sides of the larva. **B W**^⊤^**W** with **W** =[**w**_LN1_,…, **w**_LN8_] on the left and right sides. Thus each entry is 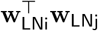, the scalar product between 2 ORNs →LN connection weight vectors **w**_LN_. **C** 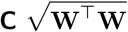, i.e., the square root of the matrices in (**B**). **D** Entries of **M** vs entries 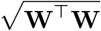, excluding the diagonal, for both sides. *r*: Pearson correlation coefficient. One-sided p-value calculated by shuffling the entries of each **w**_LN_.

**Fig. 6.**
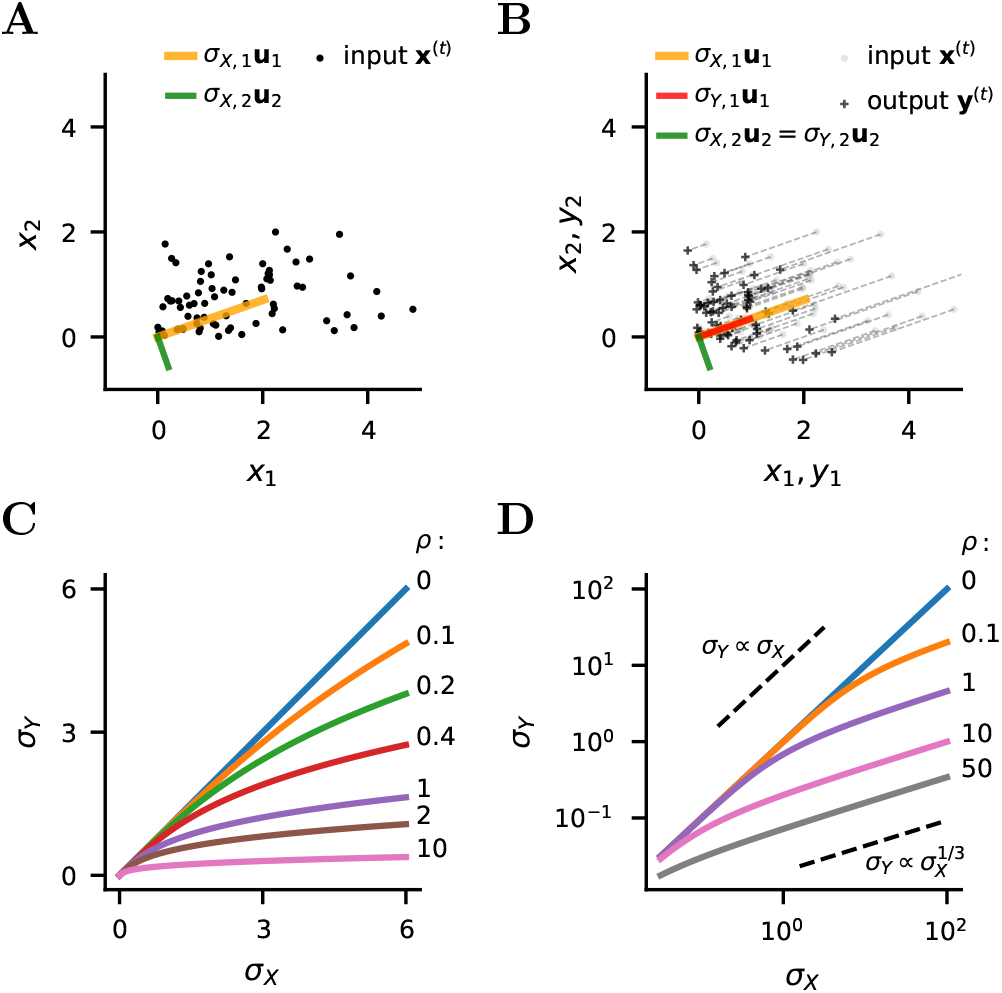
Computation in the LC. **A** Example dataset {**x**^(*t*)^} with *D* = 2 generated randomly from a zero-centered multivariate Gaussian and by removing points with negative coordinates. Depicted the PCA directions of {**x**^(*t*)^} multiplied by the s.d. of that direction. **B** Transformation from {**x**^(*t*)^} to {**y**^(*t*)^} by LC-1 (*K* = 1) with *ρ* =1. Depicted the PCA directions of {**x**^(*t*)^} and {**y**^(*t*)^} multiplied by the s.d. of that direction. **C-D** Transformation of the s.d. of PCA directions from {**x**^(*t*)^} to {**y**^(*t*)^} in the LC on linear and logarithmic axes.

**Fig. 7.**
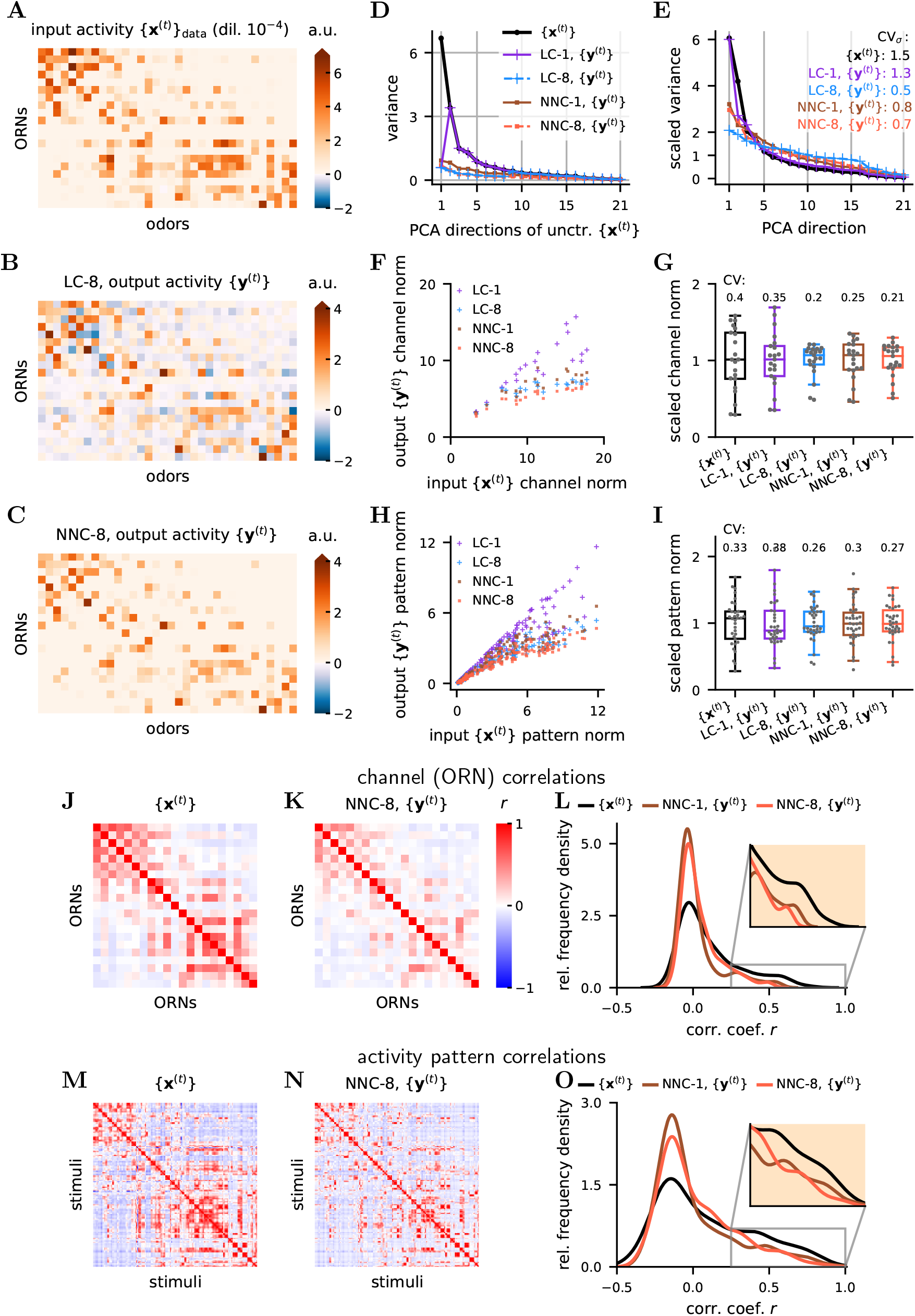
Functional consequences of LC and NNC: partial whitening, normalization, decorrelation. **A** Input (ORN soma) activity patterns {**x**^(*t*)^}_data_ for all odors at dilution 10^−4^. Instead of Δ*F*/*F*_0_ as units of activity, we use arbitrary units (a.u.), which stand for appropriate activity units at the neurons level. **B** Output {**y**^(*t*)^} for the input of (**A**) for the LC-8. **C** Same as (**B**) for NNC-8. **D** Variances of {**x**^(*t*)^}_data_ and {**y**^(*t*)^} in the principal directions of uncentered {**x**^(*t*)^}. **E** PCA variances of {**x**^(*t*)^}_data_ and {**y**^(*t*)^}, scaled by their mean. {**y**^(*t*)^} has a smaller span of variances than {**x**^(*t*)^}. See **Fig. S8** for the relation between the principal directions of {**x**^(*t*)^}_data_ and {**y**^(*t*)^}. **F** Euclidean norm of the 21 channels in output {**y**^(*t*)^} (ORN axons) vs in the input {**x**^(*t*)^}_data_ (ORN somas). **G** Box plot of the channel norms scaled by their mean, CV on top. **H** Euclidean norm of the 170 activity pattern in output {**y**^(*t*)^} vs in the input {**x**^(*t*)^}_data_ **I** Box plot of the activity patters norms (only for dilution 10^−4^) scaled by their mean, CV on top. **J** ORNs correlations in the input {**x**^(*t*)^}_data_. *K* ORNs correlations in the output {**y**^(*t*)^} of the NNC with *K* = 8. **L** Histogram for the channel correlation coefficients from (**J-K**), excluding the diagonal (n=210). **M** Activity vector (i.e., pattern) correlation in {**x**^(*t*)^}_data_ **N** Activity vector correlation in {**y**^(*t*)^} of NNC-8. **O** Histogram for the pattern correlation coefficients from (**M-N**), only for dilution 10^−4^ (n=561) (see also **Fig. S9**). *ρ* = 2 in the whole figure.

In this study, we further our understanding of LNs and their computations. We highlight the importance of minutely organized ORN - LN and LN - LN connection weights, which allows LNs to encode different significant features of input activity and dampen them in ORN axons. The transformation from the representation in ORN somas to that in ORN axons consists of a partial equalization of the PCA variances, which enables a more efficient stimulus encoding (Barlow, 1961). Indeed, this results in a decorrelation and equalization of ORNs and odor representations, which correspond to two fundamental computations in the brain: partial ZCA (zero-phase) whitening (Bell & Sejnowski, 1997; Kessy et al., 2018) and divisive normalization (Carandini & Heeger, 2012). In essence, we uncover an elegant neural circuit motif that can, via associative Hebbian plasticity, adapt to different stimuli environment and learn to extract features as well as to perform two critical computations. Thus, we present a framework that allows to quantitatively link synaptic weights in the structural data with the circuit’s function and with the circuit adaptation to input correlations, thus making a crucial step towards more integrated understanding of neural circuits.

## Results

### ORN-LN circuit

ORNs in the *Drosophila* larva carry odor information from antennas to the antennal lobe. There it is reformatted and handed over to PNs which transmit it to higher brain areas like the mushroom body and the lateral horn (Berck et al., 2016). LNs, which synapse bidirectionally with ORN axons and PN dendrites, strongly contribute to this reformatting through presynaptic and postsynaptic inhibition, as mainly shown in the adult fly (Asahina et al., 2009; Chou et al., 2010; Kim et al., 2015; Laurent, 2002; Nagel et al., 2014; Olsen et al., 2010; Olsen & Wilson, 2008).

Here, we focus on the circuit and computation presynaptic to PNs, i.e., occurring from ORN somas to ORN axons driven by LN inhibition. Specifically, we study the sub-circuit formed by all *D* = 21 ORNs and those 4 LN types (on each side of the brain) that provide direct inhibitory feedback onto the ORNs (Berck et al., 2016) (**Fig. 1A, S1**). The 4 LN types include 3 Broad Trio (BT) neurons, 2 Broad Duet (BD) neurons, 1 Keystone (KS, bilateral connections) neuron and 1 Picky 0 (P0) neuron (**Fig. S1, S2A**). This amounts to 8 ORNs - LN connections per side (3 BTs, 2 BDs, 2 KSs, and 1 P0s), and 16 on both sides.

We use the number of synapses in parallel between two neurons as a proxy of the synaptic weight *w* because synapses in the *Drosophila* larva have been found to be of similar sizes (Scheffer et al., 2020; Takemura et al., 2013) and synaptic size correlates with strength (Holderith et al., 2012). In the linear approximation, the contribution of a connection to the postsynaptic neuron activity *a_post_* is proportional to the product of *w* and the presynaptic neuron activity *a_pre_*, i.e., *a_post_* ∝ *w* · *a_pre_*.

We focus our analysis on the feedforward ORNs → LN connection weight vectors, **w**_LN_, whose *D* = 21 components are w’s corresponding to the connections from different ORNs onto the same post-synaptic LN rather than the feedback LN → ORNs. Because all the components of such a weight vector share the same post-synaptic neuron their effect on the post-synaptic activity is directly comparable, i.e. the coefficient of proportionality in *a*_LN_ ∝ ∑*_i_ w*_LN,*i*_ · *a_pre,i_* is the same. Conversely, the *w*s from one LN onto all 21 ORNs are not directly comparable among each other, because each connection affects a different postsynaptic ORN, which potentially has different electrical properties. Yet, the feedforward and feedback connection vectors are somewhat correlated (**Fig. S2**).

While Berck et al., 2016 divided the LNs into the above types based on their neuronal lineage, morphology, and qualitative connectivity, we also find that such types are innervated differently by ORNs (**Fig. 1B**). Indeed, the average correlations within LN type is higher than between LN types **w**_LN_ (**Fig. 1C,D**). Thus, for a part of our study (**Fig. 2, 3, 4A,B**) we use the 4 average 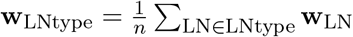, where *n* is the number of connection vectors for that LN type.

### Odor representations in ORNs are aligned with ORNs → Broad Trio connectivity weight vector

Several studies proposed that the LNs could facilitate decorrelation of the neural representation of odors (Friedrich, 2013; Friedrich & Laurent, 2001; Friedrich & Wiechert, 2014; Giridhar et al., 2011; Gschwend et al., 2015; Wanner & Friedrich, 2020). To perform such decorrelation, the circuit needs to be adapted to or “know about” the correlations in the activity patterns (Simoncelli & Olshausen, 2001). We investigated if this is the case in this olfactory circuit by testing whether the **w**_LNtype_ contain signatures of ORN activity patterns.

An ensemble of ORN activity patterns {**x**^(*t*)^}_data_ (*t* = 1,…, 170) was obtained using Ca^2+^ fluorescence imaging of ORN somas in response to a set of 34 odorants at 5 dilutions (Si et al., 2019) (**Fig. 2A**). These odorants were chosen from the components of fruits and plant leaves from the larva’s natural environment to stimulate ORNs as broadly and evenly as possible, with many odorants activating just a single ORN at the lowest concentration (i.e., the highest dilution).

Activity patterns **x**^(*t*)^ elicited by different odorants are correlated with the synaptic weight vector **w**_BT_ to a different degree (**Fig. 2B-D**), yet are such correlations statistically significant? To determine this, we first calculate the Pearson correlation coefficients *r* between the four **w**_LNtype_ and the ensemble of {**x**^(*t*)^}_data_ (**Fig. 2E**). Each **w**_LNtype_ exhibits a different “connectivity tuning curve” shape (**Fig. 2F**), **w**_BT_ being the most broadly aligned to the **x**^(*t*)^ of this stimuli set, **w**_P0_ the most sharply aligned to a few **x**^(*t*)^, and the **w**_BD_ and **w**_KS_ the most weakly aligned. To test if the **w**_LNtype_ are significantly aligned with the ensemble {**x**^(*t*)^}_data_, we compare the relative cumulative frequency (RCF) of *r* in the data with the RCFs of *r* obtained after randomly shuffling the entries of each **w**_LNtype_ (**Fig. 2G,H**). We use the maximum deviation from the mean RCF from shuffled connection vector to measure significance and find that only **w**_BT_ is significantly aligned to {**x**^(*t*)^}_data_ (**Fig. 2H,I**).

Furthermore, we find that **w**_BT_ is significantly aligned with the first PCA direction of {**x**^(*t*)^}_data_ (**Fig. S6A,B**), but none of remaining **w**_LNtype_ significantly aligned with any of the top 5 PCA directions (**Fig. 3**). We choose to compare with the top 5 (instead of 4, as the number of **w**_LNtype_) PCA directions of {**x**^(*t*)^}_data_ to cover more activity direction, thus accounting for the fact that this activity dataset does not have the same statistics of odors as the true larva environment, and likely has a different order of PCA directions. We performed PCA without centering {**x**^(*t*)^}_data_, to avoid any preprocessing on the activity data and mimic what the circuit is experiencing. The first PCA direction is thus relatively close to the mean activity direction.

Next, to test whether the connection vectors **w**_LNtype_ might be linear combinations of the PCA directions of {**x**^(*t*)^}_data_, we examine the alignment of the subspace spanned by the 4 **w**_LNtype_ and the one spanned by the top 5 PCA directions of {**x**^(*t*)^}_data_ (**Fig. S5**). We define a measure 0 ≤ Γ ≤ 4, approximately representing the number of aligned directions between these 2 subspaces (**Methods**) and find Γ ≈ 2. This value significantly deviates from the expected Γ from subspaces generated by 4 and 5 Gaussian random normal vectors in 21 dimensions (*p* < 10^−4^) and subspaces generated from the 4 connectivity vectors with shuffled entries and the 5 original activity vectors from PCA (*p* < 0.01). Approximately 1 more dimension is significantly aligned between the 2 subspaces than expected by random, supporting the results from **Fig. 3C**.

In summary, we find that **w**_BT_ is adapted to ORNs activity patterns {**x**^(*t*)^}_data_ as demonstrated by (1) the significant alignment of **w**_BT_ with individual activity patterns **x**^(*t*)^, (2) the significant alignment of **w**_BT_ with the top PCA direction of {**x**^(*t*)^}_data_, and (3) by a significantly large Γ. This supports the idea that the circuit is at least partially adapted to ORN activity patterns. This analysis fails, however, to reveal the relation between ORN activity and LNs other than BT.

### A normative and mechanistic model of the ORN-LN circuit

A detailed bottom-up modeling of the circuit requires the knowledge of the multiple unavailable physiological parameters such as ion channel distributions and neural morphologies. We therefore take here a route that circumvents these unknowns and harvests the benefits of normative approaches: similar to physics, we guess the circuit cost function, derive the governing equations, and see if their predictions agree with experiments.

Similarity-matching objective functions have been shown to be capable of extracting PCA subspaces and can be optimized by biologically plausible neural circuits with Hebbian synaptic learning rules (Pehlevan et al., 2018). Motivated by the result that the ORN-LN circuit might be adapted to at least one PCA direction of the input, we postulated a similarity-matching inspired objective function (equation (18)), such that its online optimization equations maps onto the neural dynamics of the ORN-LN circuit (equations (19), (20)) and Hebbian plasticity update rules for ORN-LN and LN-LN synapses (equation (21)). Biologically, the circuit synaptic weights could be “learned” either over evolutionary time scales, and/or during the animal lifetime.

Given a set of *T* inputs [**x**^(1)^,…, **x**^(*T*)^] = {**x**^(*t*)^}_*t*=1…*T*_ representing the activity patterns of ORN somas, the model provides us with the learned connection weights between *D* ORNs and *K* LNs: **W** = [**w**_1_,…, **w***_K_*] as well as between LNs: **M** = {*m_i,j_*}_*i,j*=1…*K*_. *m_i,i_* relates to the leak term of LN *i*. [**w**_1_,…, **w***_K_*] and **M** set the input-output relationship of the circuit and determine the activity patterns of ORN axons: {**y**^(*t*)^}_*t*=1…*T*_ and LNs: {**z**^(*t*)^}_*t*=1…*T*_. In addition to *K*, the number of LNs, the model contains only one effective parameter *ρ* characterizing the strength of the feedback inhibition.

We consider two models. First is a Linear Circuit LC-*K*, (equations (19), arising from the unconstrained objective function (18)), for which we derived an analytical solution for [**w**_1_,…, **w***_K_*], **M**, {**y**^(*t*)^}, and {**z**^(*t*)^} (**Supplementary Information**). Although linearity might be an over-simplification of the biological reality, it allows us to build up intuition. Second is a Non-Negative Circuit, NNC-*K*, (equations (20), arising from objective function (18), containing non-negativity constraints on the ORN axon and LN activity), which might be more biologically plausible. The results below for the NNC arise from numerical simulations.

### Predictions of the ORN - LN connection weight vectors

We start by analyzing the prediction of our model in terms of circuit connectivity. In the LC-*K*, the {**w***_k_*}_*k*=1…*K*_ span the subspace of the top *K* PCA directions of the input {**x**^(*t*)^} (**Supplementary Information**):

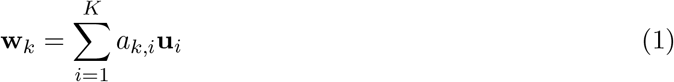

where {**u***_i_*}_*i*=1…*K*_ are the top *K* PCA directions of the dataset {**x**^(*t*)^}, {*a_i,j_*}_*i,j*=1…*K*_ are coefficients such that all **w***_k_* are linearly independent. Thus, the **w***_k_* in the LC do not necessarily correspond to specific PCA directions and are not orthogonal, and there is a degree of freedom in the {*a_i,j_*}, making the solution of the optimization not unique. Such synaptic organization assure that LNs in the LC extract the top *K* PCA subspace of the input (below). This structural prediction is tested and only partially verified in the data above (**Fig. 3**): the first PCA direction of {**x**^(*t*)^}_data_ significantly aligns with **w**_BT_, but there is no full alignment between the connectivity {**w**_LNtype_} and activity ORN principal subspaces.

Next, we study the predictions of the NNC-4 (*K* = 4 as the number of LN types). We numerically optimize the objective function (18) with {**x**^(*t*)^}_*t*=1…*T*_ = {**x**^(*t*)^}_data_ (**Fig. 2A**), *K* = 4, *ρ* = 1 and obtain {**y**^(*t*)^}, {**z**^(*t*)^}, and [**w**_1_,…, **w**_4_] (**Fig. S6C**). Intuitively, the {**w***_k_*} relate to cluster centers in soft K-means or to features in non-negative matrix factorization and the 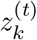 are the soft-clustering membership coefficients of **x**^(*t*)^ (below).

Three of the four **w***_k_* align significantly with the **w**_LNtype_ (BT, BD, and P0, **Fig. 4A**). This result is robust for *ρ* < 3.1 (**Fig. 4B**): all numerical optimization converge to the same {**y**^(*t*)^}, {**z**^(*t*)^}, and {**w***_k_*} for the input {**x**^(*t*)^}_data_ and given *ρ*. This can partially be attributed to the non-negativity constraint in NNC, which removes an intrinsic symmetry of the LC model. Although **w**_KS_ is the least aligned to the found **w***_k_*, NNC-5 has one **w***_k_* aligned with **w**_KS_ too (**Fig. S6H**). In summary, the ORN → LN connection weights predicted by the NNC model trained on ORN activity data {**x**^(*t*)^}_data_ largely explain the **w**_LNtype_ of the connectome. Thus, several LNs are adapted to statistical features of these ORN activity patterns.

### Emergence of LN groups in the NNC

In the connectome LNs are grouped by type and several **w**_LN_ are similar (**Fig. 1B-D, 4C**). Do LN groups naturally emerge in our model? In the LC, the {**w***_k_*}_*k*=1…*K*_ spans a *K*-dimensional subspace (given enough independent dimensions in the input {**x**^(*t*)^}). All **w***_k_* are thus different. Therefore, in the LC, LN types emerge, but no similar LNs. In the NNC with small *ρ*, however, the objective function (18) leads to the symmetric non-negative matrix factorization (SNMF) objective function between (**x**^(*t*)^} and {**z**^(*t*)^} (**Supplementary Information**), which corresponds to a soft clustering of **x**^(*t*)^ by **z**^(*t*)^. Thus, each component in **z**^(*t*)^ discovers and encodes the presence of a sparse feature of **x**^(*t*)^ (Pehlevan & Chklovskii, 2015). In that case, when the number of significant sparse features in {**x**^(*t*)^} is smaller than *K*, several components of **z**^(*t*)^ (i.e., LNs) encode a similar feature. Our simulations for NNC-8 (*K* = 8 as the number of LNs on each side of the larva) with {**x**^(*t*)^}_data_ and *ρ* = 0.1 indeed give rise to groups of similar **w***_k_* (**Fig. 4D**). Conversely, for larger p, the **w***_k_* become more decorrelated (**Fig. 4D**, *ρ* = 10). To study how the resemblance of the **w***_k_* changes with *ρ*, we calculated the average rectified correlation coefficient 〈*r*_+_〉 between all the **w***_k_* for different *ρ* (**Fig. 4D,E**). At *ρ* = 0.35, 〈*r*_+_〉 of the NNC-8 matched that of the connectome. This value of *ρ* should not, however, be interpreted as a “true” value for the actual biological circuit, because the true ORN activity patterns {**x**^(*t*)^} that the larva experienced is unknown - in fact changing {**x**^(*t*)^} and *ρ* are two independent means of influencing the model circuit synaptic weights. In summary, within reasonable parameter ranges, the NNC reproduces yet another property of the biological circuit: the emergence of LNs that can be grouped by type.

### Relation between LN-LN and feedforward ORNs → LN connection weights

The ORN - LN circuit also contains inhibitory reciprocal LN - LN connections (**M** = {*m*_LNi, LNj_}, **Fig. 5A**) whose role is not fully understood. Our model predicts that **M** and **W** = [**w**_1_,…, **w***_K_*] are related thus (**Supplementary Information**):

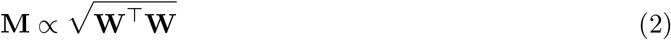

Where ⊤ is the matrix transpose. This relationship is exact for the LC and approximate for the NNC. First, it predicts that the matrix **M** is symmetric, i.e., that the synaptic weight of LN*_i_* → LN*_j_* is equal to that of LN*_j_* → LN*_i_*. This is indeed approximately true in the connectome, except for the P0, which inhibits KS, but is not strongly inhibited by them (**Fig. 5A**). Second, as predicted by the relationship (2), we find, in the connectome, a significant correlation between the entries of **M** and 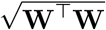 for the left and right sides of the larva (excluding the diagonal entries, since the connectome does not provide the values corresponding to the diagonal of **M** of the model circuit) (**Fig. 5**). This suggests that the ORN-LN and LN-LN connections are meticulously co-organized to perform the circuit’s function. Intuitively, LN-LN interaction could be interpreted as LNs competing with each other for activation. During circuit learning, without LN-LN connections, all LNs would learn the same most significant direction of the input data. Thus, these lateral connections ensure that LNs span more than a single direction of the ORN activity space. After learning, LN-LN connections constitute an essential part of the computation (below, **Fig. S11**).

In summary, the NNC model accurately predicts several key features of the connectome: the **w**_LNtype_ connection weights, the emergence of LN groups, and the relationship between ORNs → LN and LN - LN connections weights.

### Computation in the LC: partial equalization of PCA variances in ORN axons and extraction of principal subspace by LNs

Next, we examine the computation performed by the LC model. The computation is implemented dynamically through the ORN - LN loop and converges exponentially to a steady state (equation (19)). Given inputs {**x**^(*t*)^}, we consider the twofold output of the circuit: the converged representations in ORN axons {**y**^(*t*)^} and in LNs {**z**^(*t*)^}, both transmitted downstream. Although LNs are usually thought of only performing local computations, here LNs also project to several types of neuron like uni- and multi-glomerular PNs (Berck et al., 2016). Because the circuit is adapted to its input {**x**^(*t*)^}, the transformations from **x**^(*t*)^ to **y**^(*t*)^ and **z**^(*t*)^ are related to the statistics of {**x**^(*t*)^} and are naturally expressed using the PCA directions {**u***_i_*} and variances 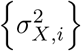 (*i* = 1,…, *D*) of uncentered {**x**^(*t*)^}. Formally, given the autocorrelation matrix 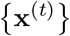 and **u***_i_* are the eigenvalues and eigenvectors of **Σ***_X_*, respectively (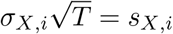 is also the i^th^ singular value of {**x**^(*t*)^}), **U** = [**u**_1_,…, **u***_D_*], and **Λ***_X_* = diag(*σ*_*X*,1_,…, *σ_X,D_*). We write the odor representations in ORN somas in this basis and find (**Supplementary Information**):

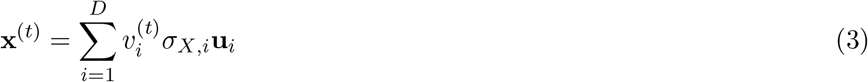

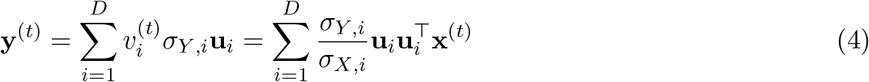

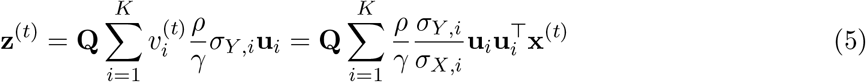

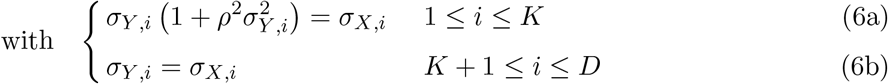

where 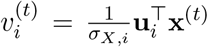 are the coefficients of **x**^(*t*)^ in the orthogonal basis {*σ_X,i_**u**_i_*} and **Q** is a (*K* × *K*) orthonormal (rotation) matrix and is a degree of freedom of the optimization.

On the dataset level, we find 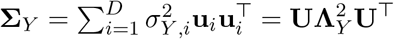 where **Λ***_Y_* = diag(*σ*_*Y*,1_,…, *σ_Y,D_*). Thus, the activity patterns in ORN axons {**y**^(*t*)^} have the same principal directions {**u***_i_*} as {**x**^(*t*)^} but with modified PCA variances (portrayed in **Fig. 6A,B** with *D* = 2 and *K* = 1). The variances of the last *D* – *K* PCA directions of {**x**^(*t*)^} remain unaltered in {**y**^(*t*)^}, whereas the variances of top *K* directions (as the number of LNs) are diminished according to equation (6a) (**Fig. 6C,D**), because LNs ({**z**^(*t*)^}) encode (a rotated version of) the top *K* principal subspace of {**x**^(*t*)^} (equation (5)) and inhibit it in the ORN axons ({**y**^(*t*)^}). From the top *K* principal directions, those with relatively large variances are shrunken with a cubic root 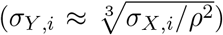, whereas those with relatively small variance remain virtually unchanged. Indeed, in the latter case, LNs are weakly activated and inhibition is almost inexistent.

For a LC with the same number of LNs as ORNs (i.e., *D* = *K*), this computation leads to a flatter spectrum of 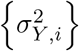 relatively to the one of 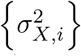, which can be quantified by the coefficient of variation, CV*_σ_* (**Supplementary Information**). Although for *K* < *D* only the top *K* principal direction are shrunken, in most cases it also leads to a decrease of CV*_σ_* (see below).

This computation is a partial (Zero-phase) ZCA-whitening. By definition, a multivariate random variable **A** is white if its autocovariance matrix is proportional to the identity matrix: **E** [(**A** – **E**[**A**]) (**A** – **E**[**A**])^⊤^] ∝ **I**, which implies that all the PCA variances (i.e., eigenvalues of the autocovariance matrix) are equal. For the LC, the CV*_σ_* of 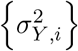 is smaller than the CV*_σ_* of 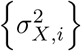 (see also **Fig. 7E** below). Although these are formally the variances of the PCA on uncentered data, because the mean of {**x**^(*t*)^}_data_ is close to **0**, flattering the spectrum of 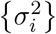 causes the flattening of the spectrum of the eigenvalues of the autocovariance matrix too, leading to partial whitening. Finally, since the principal directions of {**y**^(*t*)^} and {**x**^(*t*)^} are the same, the transformation contains no rotation and is thus “zero-phase”, as ZCA-whitening.

### LC and NNC computation on the ORN activity dataset

Finally, to elucidate the computation of this circuit on odor representations, we study the computation of the LC and the NNC on {**x**^(*t*)^}_data_. We set the parameter regulating the strength of the inhibition *ρ* = 2 to distinctly portray the input-output transformation. Given the input of ORN activities {**x**^(*t*)^}_data_, we calculate {**y**^(*t*)^} and {**z**^(*t*)^} with *K* = 1 and *K* = 8 using the analytical formula for the LC and by optimizing the objective function (18) for the NNC.

In the LC, LNs encode the top *K* principal subspace of {**x**^(*t*)^} (above, **Fig. S7B**). In the NNC, the computation in LNs approximates SNMF for small *ρ* (**Supplementary Information**) which performs soft clustering and sparse feature discovery (Pehlevan & Chklovskii, 2015). LNs thus encode features of the odor representations in ORN (**Fig. S7C-G**), that are transmitted to downstream brain areas.

Next we show that in LC and NNC the transformation from {**x**^(*t*)^} to {**y**^(*t*)^} is a partial ZCA-whitening and a divisive normalization as reflected in the partial equalization of the PCA variances (**Fig. 7E**), the decrease of channel (i.e., ORN) and pattern (i.e., neural representations of odors) correlations (**Fig. 7J-O, S9**), and the lack of rotation of the output (**Fig. S8E**). **Fig. 7A-C** shows the activity in ORN somas and the computed activity in ORN axons for LC-8 and NNC-8. The LC produces strongly negative values in {**y**^(*t*)^}, which might not be biologically plausible. We next compared the spectrum of 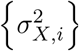 and 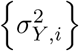, since this characterizes whitening and the computation in LC affects this aspect (**Fig. 7D**). As expected, in the LC only the top *K* principal directions of the input are dampened. For the NNC, however, we find that all directions are dampened, even for *K* = 1. This can be attributed to the non-negativity constraint on the output {**y**^(*t*)^} and {**z**^(*t*)^} in NNC, which potentially affects all stimuli directions. We find a flattening of 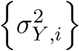 spectrum both in LN and NNC as seen in the smaller CV*_σ_* (**Fig. 7E**) demonstrating that {**y**^(*t*)^} is more white that {**x**^(*t*)^}. Changing the number of LNs does not affect the NNC as much as the LC. However, changing *ρ* greatly influences the strength of the dampening (**Fig. S10**). Although in the LC the principal directions of {**x**^(*t*)^} and {**y**^(*t*)^} remain the same, their order changes, because only a fraction of them are shrunken (**Fig. S8A,B**). For the NNC, however, there is only a slight mixing between principal directions of similar strength, but their order mainly remains (**Fig. S8C,D**).

As expected from a flatter {*σ_Y,i_*}, we observe that channels and patterns are more decorrelated in the output {**y**^(*t*)^} in the NNC (**Fig. 7J-O**) and in the LC (**Fig. S9**) than in the input, which is coherent with partial whitening. The strength of decorrelation increases with *ρ* (**Fig. S10**).

Next, we study the effect of the circuit computation on channel and pattern activity Euclidean norms, which reflect the total channel and total pattern activity. We find that both LC and NNC dampen the channels with strong norms and leave the weaker channels largely unaffected, thus decreasing the CV of channel norms (**Fig. 7F,G**). This allows the information to be more evenly distributed among channels, an important property of efficient coding. Similarly, the circuit partially equalizes the norms of activity patterns (**Fig. 7H,I**). This slightly removes the concentration information from the signal. These effects are similar to a divisive normalization-type computation, also reported in *Drosophila* (Carandini & Heeger, 2012; Olsen et al., 2010).

Finally, we aim at better understanding the role of LN-LN connections. We study the computations performed by the converged LC and NNC, with the off-diagonal elements in **M** set to 0 (**Fig. S11**). We find that this manipulation mixes the output principal direction in relation to the input and also increases the total level of inhibition. Thus, LN-LN connection helps to reduce the amount of rotation in the neural representation, regulate the amount of inhibition, and maintain the predicted computation.

In summary, the analysis of the LC and NNC predicts that the ORN-LN circuit performs the following computation on the odor representation in ORNs: it most strongly dampens the most prominent directions of the input dataset and thus flatten the PCA variance spectrum. This results in an output in ORN axons that is more white, decorrelated, and more equalized channels and patterns. This allows a more efficient neural representation and improves odor discrimination downstream.

## Discussion

Combining the *Drosophila* larva olfactory circuit connectome, ORN activity data, and a new normative model, we advance the understanding of sensory computation and adaptation, quantitatively link ORN activity statistics, functional data and connectome, and make testable predictions. Our work uncovers and characterizes a simple and potent neural circuit architecture capable of adaptive data preprocessing and feature extraction, which, as an independent computational unit, could arise in other brain areas and be useful for machine learning and signal processing. Finally, our normative approach provides a general framework to understand circuit computation (Bahroun et al., 2019; Golkar et al., 2020) and could be applied to more connectomes (Eichler et al., 2017; Scheffer et al., 2020).

### Circuit computation, partial ZCA-whitening, and divisive normalization

We propose that the circuit’s effect on neural odor representation in ORNs correspond to partial ZCA-whitening and divisive normalization (DN) (**Fig. 6, 7**). Such computations, which reduce correlations originating from the sensory system and the environment, have appeared in efficient coding and redundancy reduction theories (Atick & Redlich, 1992; Barlow, 1961; Carandini & Heeger, 2012; Linsker, 1988; Plumbley, 1993; Simoncelli & Olshausen, 2001). Partial whitening is indeed a solution for mutual information maximization in the presence of input noise (Atick & Redlich, 1990). In this circuit too, we suggest that a pure whitening transformation might not be desirable, as it could lead to noise amplification. Thus, keeping low-variance signal directions of the input unchanged and damping larger ones might accord with mutual information maximization. Our conclusions are in line with reports of pattern decorrelation and/or whitening in the olfactory system in zebrafish (Friedrich, 2013; Friedrich & Laurent, 2001; Friedrich & Wiechert, 2014; Wanner & Friedrich, 2020) and mice (Giridhar et al., 2011; Gschwend et al., 2015).

Infinitely many whitening transformations exist - indeed, a rotated white signal remains white. ZCA-whitening, where the output is not rotated relatively to the input, might be advantageous over other flavors of whitening because it is the optimal whitening transform that minimizes the distance between the original and the whitened signal (Kessy et al., 2018). Since inputs (i.e., spike rates) are non-negative, this property of ZCA-whitening will reduce the amount of negative deviations and lessen the distortion of the computation that arises from the non-negative constraint on neural activity.

On the other hand, the computation in our model also resembles DN, a ubiquitous computation in the brain (Carandini & Heeger, 2012) which was suggested for the analogous circuit in the adult *Drosophila* (Olsen et al., 2010; Olsen & Wilson, 2008). In its simplest form, DN is defined as 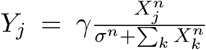, where *Y_j_* is the response of the neuron *j, X_i_* is the driving input of the neuron *i*, and *γ, σ*, and *n* are positive parameters. DN captures two effects of neuronal and circuit computation: (1) the saturation of a neural response with increasing input up to a maximum spiking rate *γ*, which mainly arises from neuron’s biophysical properties; (2) dampening of the response of a given neuron when other neurons also receive input, usually originating from lateral inhibition (but see Sato et al., 2016). In our model, aspect (1) of DN is absent, but could readily be implemented with a saturating non-linearity. However, signatures of (2) are especially apparent in the saturation of the pattern output norm for increasing input norm (**Fig. 7H**). This saturation occurs because inputs with higher norms correspond to inputs at higher odor concentrations and with a higher number of active ORNs. Because such input directions are more statistically significant in our dataset, these stimuli that are more strongly dampened by LNs (which encode those directions) than those with few ORNs active. Thus, our model presents a possible linear implementation of a crucial aspect of DN, which in itself is a nonlinear operation.

The basic form of DN equalizes the channels and performs channel decorrelation, but not pattern decorrelation (Friedrich & Wiechert, 2014; Olsen et al., 2010; Wanner & Friedrich, 2020), which appears in our model. However, a modified version of DN, which includes different coefficients for the driving inputs in the denominator (Westrick et al., 2016), performs pattern decorrelation too, as seen in our circuit. The proposed neural implementations of DN usually require a multiplication by the feedback (Heeger, 1992; Westrick et al., 2016), which might not be as biologically realistic as our circuit implementation.

Several neural architectures similar to ours have been proposed to learn to decorrelate channels, perform DN, or learn sparse representations in an unsupervised manner (Atick & Redlich, 1993; King et al., 2013; Koulakov & Rinberg, 2011; Olshausen & Field, 1997; Pehlevan & Chklovskii, 2015, 2016; Westrick et al., 2016; Wick et al., 2010; M. Zhu & Rozell, 2015). These studies, however, either do not have an objective function, or have a different circuit architecture or synaptic learning rules.

### Roles of LNs

LNs form a significant part of the neural populations in the brain, have multiple crucial computational functions, and have extremely diverse morphologies and excitabilities (Chou et al., 2010; Hattori et al., 2017). We propose a dual role for LNs in this olfactory circuit: altering the odor representation in ORNs and extracting ORN activity features, which can be used downstream (Berck et al., 2016). In the olfactory system of *Drosophila* and zebrafish, LNs perform multiple roles like gain control, normalization of odor representations, pattern and channel decorrelation (Friedrich, 2013; Friedrich & Wiechert, 2014; Olsen et al., 2010; Olsen & Wilson, 2008; Wanner & Friedrich, 2020; P. Zhu et al., 2013), roles that are in line with our results. Also, in *Drosophila* the LN population expands the temporal bandwidth of synaptic transmission and temporally tune PN responses (Kim et al., 2015; Nagel et al., 2014; Nagel & Wilson, 2016), which was not addressed here.

In topographically organized circuits such as visual periphery or auditory cortex, several LN types uniformly tile the topographic space and each LN type has its own role and selectivity (e.g., in the retina (Masland, 2012)). In non-topographically organized networks, however, the organization and selectivity of LNs is still a matter of research and controversy (Chou et al., 2010; Hong & Wilson, 2015). We have included 4 LN types in the studied subcircuit (**Fig. 1**). Several LN types contains multiple copies of LNs, with similar connection weights, and thus presumably similar roles. In the LC model, the *K* LNs span a *K*-dimensional subspace of activity, thus each LN has a different connectivity and would form a type of its own. In the NNC model, large *ρ* lead to different LNs, whereas smaller *ρ* lead to the formation of LN groups (**Fig. 4C-E**). Thus based on our study and the different connectivity patterns of LNs in the connectome (Berck et al., 2016), we suggest that in the *Drosophila* larva LN types extract different features of ORN activity and are thus differently activated in response to different input directions (and glomeruli) and also different ORNs are differently inhibited by different LNs. This seems at odds with the results of Hong and Wilson, 2015 who found that the activation of the LN population appears invariant to odor identity. However, the latter study imaged several LNs simultaneously and thus might have missed the selectivity of individual LNs.

What are the features being extracted by LNs? The Broad Trio, whose connection weight vector aligns to the first PCA direction of ORN activity and to a **w** of the NNC model (**Fig. 3, 4A,B**), could potentially encode the mean ORN activity, and thus be related to the global odor concentration (Asahina et al., 2009). Other LNs, whose connectivity aligns with the **w** of the NNC model, might encode features of odors, like aromatic vs long carbon chain (Si et al., 2019), or specific information influencing larva behavior (Berck et al., 2016). What is the function of multiple “copies” of LNs within each type? Firstly, LNs might differentiate further as the larva grows, and as the circuit continues learning. Secondly, several LNs might help expand the dynamical range of a single LN.

The connectome reveals that the circuit also includes LN-LN connections, which arise naturally in our approach. We suggest that LN-LN connections constitute a crucial part of learning and LN differentiation, as well as performing partial ZCA-whitening and normalization. Our model also correctly predicted how LN-LN connections co-organize with the ORN-LN connections (**Fig. 5**). To our knowledge, the role of LN-LN connections and their relationship to ORN-LN connections has not been addressed previously in such circuits.

In summary, our study highlights the significance of the different ORN-LN and LN-LN connection strengths and argues that LNs are minutely selective and organized to extract features and render the representation of odors more efficient.

### Learning and ORN activity statistics

Using ORN activity dataset (Si et al., 2019), our NNC model could predict to a large extent the connection weight vectors found in the connectome (**Fig. 4A-B**). This suggests that the circuit is adapted to ORN activity patterns (**Fig. 2, 3, 4**). How could the connectivity prediction be successful, when the ORN activity dataset was mainly chosen to uniformly and broadly activate all ORNs and not to match the true larva odor environment, in terms of odor identity, frequency, and intensity? One possibility is that, given an ORN activity dataset large enough, certain generic correlations between ORNs always appear, giving rise to the same robust features in the connectivity. These correlations could be caused by intrinsic chemical properties of ORN receptors. Moreover, the exact odor statistics would also alter the connection weights, but to a lesser extent than the former effect. Thus, given an activity dataset closely mimicking the larva natural odor environment, the model predictions of the connectome might further improve.

Are those synaptic weights learned during the animal lifetime or are they encoded genetically, i.e., “learned” over an evolutionary time span? A genetic origin is undoubtedly present, given that several LNs types (e.g., Keystone and Picky) differ by their connectivity to specific neurons outside the studied circuit and seem to be linked to different hard-wired animal behaviors (Berck et al., 2016). Additionally, several studies reveal that glomeruli sizes (and thus ORN-LN or ORN-PN synaptic weights) or activity vary depending on the environment where the *Drosophila* grows up (Arenas et al., 2012; Das et al., 2011; Devaud et al., 2001; Sachse et al., 2007; Sudhakaran et al., 2012). This feature would equip the circuit with a potent mechanism to adapt to evolving natural environment. Additionally, synaptic count and innervation variability arises for *Drosophila* brought up in similar environments (Chou et al., 2010; Tobin et al., 2017), indicating the potential imprecision of the development and/or learning. Resolving connectomes of larva raised in different odor environments, probing the synaptic plasticity present in the network, and recording ORN responses to the full ensemble of odors present in its environment would help clarify the influence of learning and of genetics.

In conclusion, our work uncovers a canonical circuit model that could robustly adapt to different environments in an unsupervised manner, while maintaining the critical computations of partial whitening, normalization, and feature extraction. Our comprehensive normative approach, which contains only one effective parameter, predicted the structural organization based on input activity, and found in the connectome the signatures of circuit function and adaptation to ORN pattern statistics. Such an approach could provide important insights into more complicated adaptive neural circuits, whose structural and activity data is becoming available.

## Methods

### ORN activity

We use the average maximal Ca^2+^ Δ*F*/*F*_0_ responses among trials for the activity data as in Si et al., 2019. For the ORN 85c in response to 2-heptanone, and for the ORN 22c in response to methyl salicylate, we only have responses to dilutions ≤ 10^−7^. Because the ORN responses are very similar for dilutions 10^−7^ and 10^−8^ and are already saturated (for this cell we have responses down to dilutions of 10^−11^), we set the missing response for dilutions 10^−6^, 10^−5^ and 10^−4^ as the response for 10^−7^.

### RCF distribution of correlation coefficient and significance testing

Given a vector 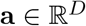, we define the mean 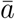, the centered vector **a***_c_*, and the centered normalized vector 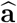:

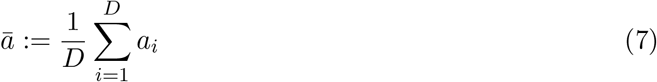

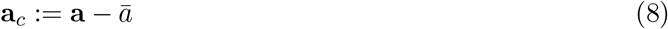

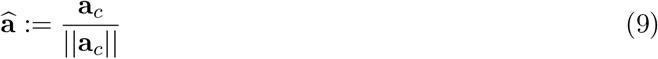

We call 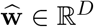 the centered and normalized ORNs → LN synaptic weight vector **w**. Similarly, we define 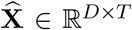 the centered and normalized ORN activity **X**_data_ = [**x**^(1)^,…,**x**^(*T*)^], where each column vector is centered and normalized.

Each row of the matrix of correlation coefficients depicted in **Fig. 2E** is given by 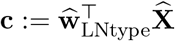. **c** is used to calculate the true relative cumulative frequency (RCF) of correlation coefficients in **Fig. 2G**: 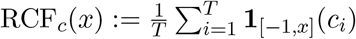, where **1***_A_*(*y*) is the indicator function of a given set *A*.

We define the random variables **w**′, **c**′ and *RCF*′. **w**′ is generated by shuffling the entries of a connectivity vector 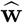:

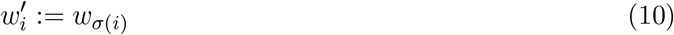

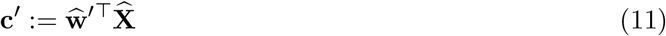

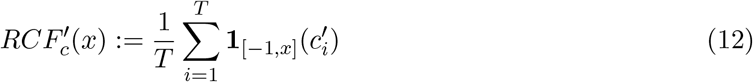

Where *σ*(*i*) is a random permutation operator. We define 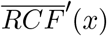 (**Fig. 2G**, black line) as the mean *RCF*′(*x*) arising from all RCFs that come from shuffled 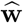. Next, we define, the maximum negative deviation *δ*′ random variable as:

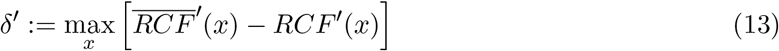

Finally, we define p-value = Pr(*δ*′ ≥ *δ_true_*). The p-value is thus the proportion of RCFs generated with random shuffling of entries of 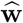 that deviate from the mean RCF more than the true RCF.

Numerically, these calculations were done by binning the RCF function into 0.02 bins and generating 10000 instances of shuffled 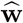.

### Number of aligned dimensions between two subspaces

Given a Hilbert space of dimension *D*, we define Ω - a measure of dissimilarity between 2 subspaces **S***_A_* and **S***_B_* generated by the matrices of linearly independent *K_A_* and *K_B_* column vectors: 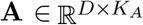 and 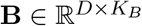:

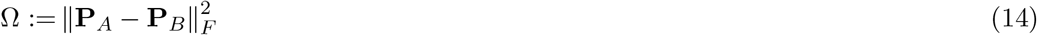

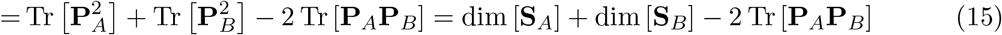

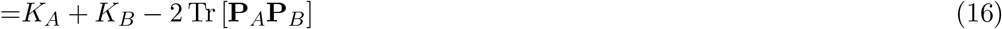

Where **P***_A_*, 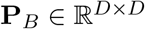 are the projectors onto the subspaces *S_A_* and *S_B_*, respectively, *F* stands for the Frobenius norm, Tr is the matrix trace, and *K_X_* = dim(**S***_X_*) is the dimensionality of a subspace *S_X_*. We assume *K_A_* + *K_B_* ≤ *D*. We have that |*K_A_* – *K_B_*| ≤ Ω ≤ *K_A_* + *K_B_*. The projection matrix can be obtained thus **P***_A_* = **A** (**A**^⊤^**A**)^−1^ **A**^⊤^, or via QR factorization: **QR** = **A, P***_A_* = **QQ**^⊤^.

Intuitively, for two very similar subspaces, the projection **P***_A_υ* of an arbitrary vector *υ* onto *S_A_* will be very similar to the projection **P***_B_υ* vector *υ* onto *S_B_*, thus **P***_A_υ* ≈ **P***_B_υ* and Ω will be small. Conversely, if the subspaces are very different, the projections **P***_A_υ* and **P***_B_υ* will also be different and Ω will be large.

We now define the more intuitive measure:

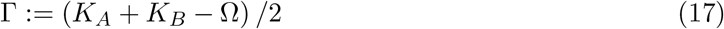

which is a proxy of the number of aligned dimensions in the two subspaces. Indeed 0 ≤ Γ ≤ min(*K_A_, K_B_*). For 2 perpendicular subspaces, Γ = 0 and for 2 fully aligned subspaces Γ = min(*K_A_, K_B_*).

In the main text we have **A** = [**w**_BT_, **w**_BD_, **w**_KS_, **w**_P0_] and **B** is the matrix with the top 5 PCA loading vectors of {**x**^(*t*)^} as columns, *K_A_* = dim [**S***_A_*] =4, *K_B_* = dim [**S***_B_*] = 5 and *D* = 21.

### Objective function for the ORN-LN circuit

We choose a normative-theoretical approach to study the ORN-LN circuit. It has the advantage of providing analytical expressions describing different aspects of the computation and the circuit architecture. Studying the circuit’s computation is then equivalent to studying the optimum of a cost function.

We first define the following variables: an input **X** = [**x**^(1)^,…, **x**^(*T*)^] of *T* samples, and outputs **Y** = [**y**^(1)^,…, **x**^(*T*)^], **Z** = [**z**^(1)^,…, **x**^(*T*)^]. **x**^(*t*)^ and **y**^(*t*)^ are *D*-dimensional vectors, whereas **z**^(*t*)^ are *K*-dimensional. **x**^(*t*)^, **y**^(*t*)^, and **z**^(*t*)^ represent the activity of ORN somas (i.e., the inputs), ORN axons and *K* LNs, respectively. We postulate the following similarity-based objective function (e.g., Pehlevan et al., 2018), which links the steady state activity of the outputs to that of the input:

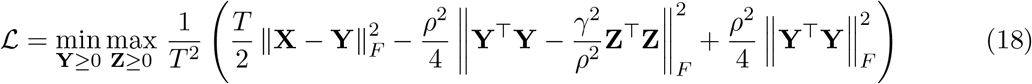

Intuitively this objective function drives the activity of the ORN axons **Y** to be close to the activity of ORN somas **X** through the term 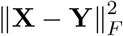, it aligns the similarity between the activity of ORN axons and LNs through the term 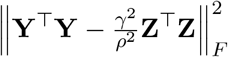, and finally puts a 4^th^ order penalty on the norm of **Y** through the term 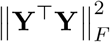. *ρ* and *γ* are two parameters. Scaling *ρ* is related to the strength of the dampening in **Y** and affects both the optima of **Y** and **Z**. Changing *γ* only scales **Z**, without affecting **Y**. Since *γ* does not fundamentally change the computation, we set *γ* = 1 in the whole paper.

We consider two objective functions. One without the non-negativity constraints on **Y** and **Z**, representing the Linear Circuit (LC) model, and one with the non-negativity constrains as in equation (18), representing the Non-Negative Circuit (NNC) model. Non-negativity constraints account for the fact that neural activity is usually non-negative, or at least not symmetric in the negative and positive directions.

In order to map the objective function to a neural circuit (**Supplementary Information**), we first introduce two auxiliary matrices 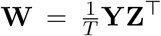 and 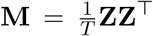, which naturally map onto ORNs - LNs and LNs - LNs synaptic weights, respectively. The objective function is thus optimized over the variables **Y, Z, W**, and **M**. We then consider the objective function in the “online setting”. In this situation one **x**^(*t*)^ is presented at a time, the optimal **y**^(*t*)^ and **z**^(*t*)^ are found with the current **W** and **M**, and subsequently the **W** and **M** are updated. The optimal **y**^(*t*)^ and **z**^(*t*)^ are found with gradient descent/ascent equations, which also correspond to the ORN-LN neural dynamics equations ((19) for the LC or (20) for the NNC). The gradient descent/ascent steps on **W** and **M** correspond to the Hebbian learning update rules equation (21).

### Circuit neural dynamics

When optimized online, the objective function (18) without the non-negativity constraints gives rise to the following differential equations describing the LC, whose steady state solutions correspond to the optima for **y**^(*t*)^ and **z**^(*t*)^ (**Supplementary Information**). These equations naturally map onto the ORN-LN neural circuit dynamics (dropping the sample index *t* for simplicity of notation):

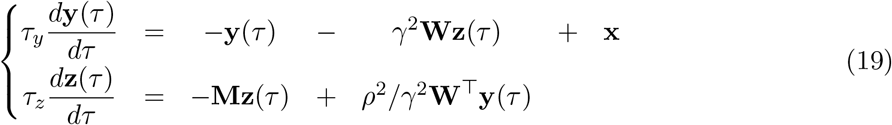

Where **x, y** and **z** are *D, D*, and K-dimensional vectors, and represent the activity (e.g., spiking rate) of the ORN somas, ORN axons, and LNs, respectively. *τ_y_* and *τ_z_* are neural time constants, *τ* is the local time evolution (not to be confused with the *t* sample index). The elements of the *D* × *K* matrices *ρ*^2^/*γ*^2^**W** and *γ*^2^**W** contain the synaptic weights of the feedforward ORNs → LN and feedback LN → ORNs connections, respectively. Thus, the feedforward connection vectors are proportional to the feedback vectors, with a scaling factor *ρ*^2^/*γ*^4^. This assumption is reasonable considering the connectivity data (**Fig. S1, S2B**). Off-diagonal elements of the *K* × *K* matrix **M** contain the weights of LN - LN inhibitory connections, whereas the diagonal elements are related to the LNs leak. In the absence of LN activity and at steady state, the equations satisfy **y** = **x**, i.e., ORN soma and axonal activities are identical. In the absence of input (i.e., **x** = 0) both **y** and **z** decay exponentially to **0**, because of the terms –**y**(*τ*) and –*M_i,iz_i__*(*τ*), respectively. In summary, these equations effectively model the ORN-LN circuit dynamics by implementing that (1) the ORN axonal activity is driven by the input in ORN somas **x** and inhibited by the feedback from the LNs thought the term –*γ*^2^**Wz**(*τ*) and (2) LN activity is driven by the activity in ORN axonal terminals by *ρ*^2^/*γ*^2^**W**^⊤^**y**(*τ*) and inhibited by LNs through the term –**Mz**(*τ*). *ρ* and *γ* are two parameters. In fact, a general system of differential equations describing this circuit architecture can be reduced to having just two parameters (**Supplementary Information**). Scaling *ρ* affects both the steady state solution of **y** and **z**, whereas scaling *γ* only scales **z**. Note that changing *ρ* in the objective function, will also give rise to different optimal **W** and **M**.

When optimized online, the objective function (18) with the non-negativity constraints gives rise to the following equations describing the NNC:

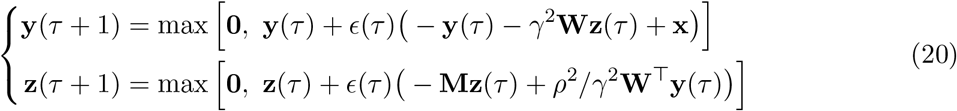

Where *ϵ*(*τ*) is the step size parameter and the max is performed component wise. Here *τ* is a discrete time variable. These equations can be seen as the equivalent to equations (19), but also satisfying constraints on the activity, such as *y_i_*(*τ*) ≥ 0, *z_i_*(*τ*) ≥ 0, ∀*τ, i*. Such constraints are implemented by formulating circuit dynamics in discrete time and using a projected gradient descent.

We call LC-*K* the linear circuit implemented by (19) and NNC-*K* the non-negative circuit implemented by (20), with *K* LNs. The actual biological circuit might exhibit a behavior somewhere between the LC and NNC. For the circuit studied here, we have *D* = 21 (number of ORNs), and *K* = 8 (number of LNs on one side of the larva) or *K* = 4 (number of LN types) or *K* = 1 (to build intuition).

### Mathematical description of synaptic plasticity

When the objective function (18) is optimized online, we obtain the following updates for **W** and **M** after each presentation of a sample **x**^(*t*)^ and convergence to optimal **y**^(*t*)^ and **z**^(*t*)^:

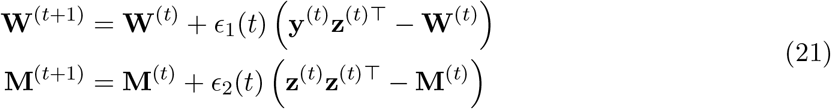

Where *ϵ_i_*(*t*) are learning rates. We assume that the ORN soma activation **x**^(*t*)^ in present long enough so that **y**^(*t*)^(*τ*) and **z**^(*t*)^(*τ*) reach steady state values. These equations represent Hebbian plasticity in **W** and **M**, which is a form of correlative unsupervised learning. This is justified by (1) the adaptation of the connectivity to statistics of the ORN activity found in our data, (2) the presence of activity-dependent plasticity in *Drosophila* (Arenas et al., 2012; Das et al., 2011; Devaud et al., 2001; Sachse et al., 2007; Sudhakaran et al., 2012), and (3) that glomeruli activity is best explained with glomerulus-glomerulus inhibitory connectivity that is proportional to the correlation between glomeruli (Linster et al., 2005). These equations (21) set the diagonal values of **M** by analogy to the off-diagonal ones.

With appropriate learning rates, these synaptic update rules lead to:

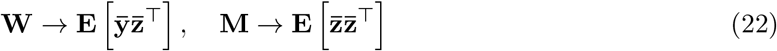

Such **W** and **M** could potentially arise either over evolutionary time scales, or during the animal lifetime. In summary, based on the postulated objective function (18), we derived neural dynamics equation (equations (19) for LC, (20) for NNC) which map onto the ORN-LN circuit and biologically plausible Hebbian synaptic plasticity rules (equations (21)). This fully specifies the circuit, its synaptic weights, and its input-output relationship.

### Numerical simulation of the LC offline

For the LC, we have the theoretical solution, so numerical simulations are not necessary to obtain **Y**. Also, there is a degeneracy in the solutions of **Z, W**, and **M**. However, to confirm the theoretical results, we did simulate the LC too. For that, we used the following equation, where the cost function depends on **Z** only (**Supplementary Information**, equation (S49), with *γ* = 1):

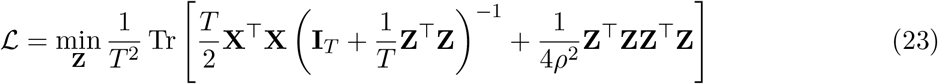

We used an algorithm similar to Kuang et al., 2012.

#### Algorithm 1 Finding the minimum of 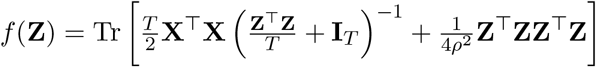

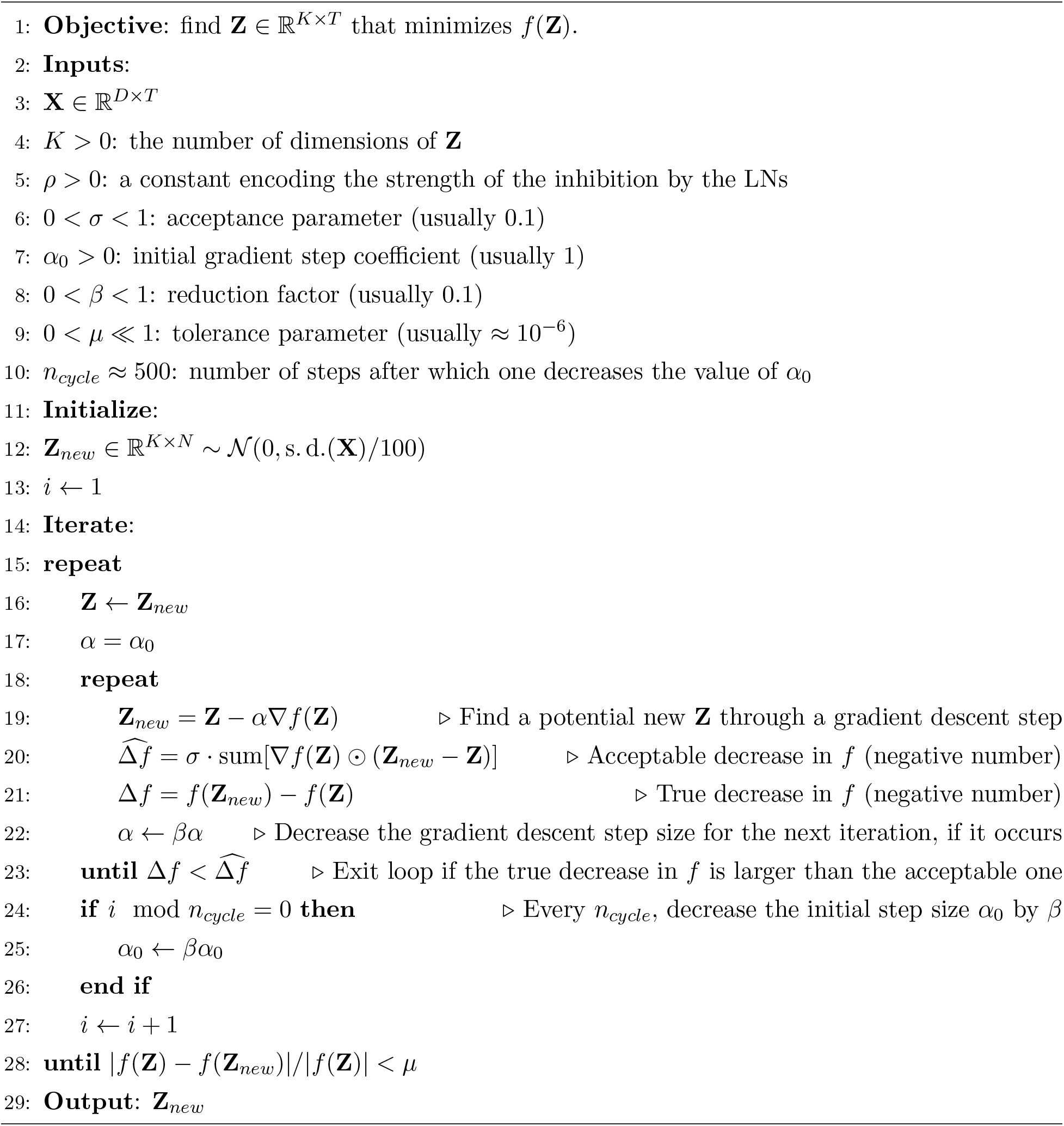

Where ʘ is an element-wise multiplication and the “sum” adds all the elements of the matrix. In the inner repeat loop of the algorithm, it can happen that because of limited numerical precision, no α is small enough to make a decrease in *f* (i.e., satisfy the condition 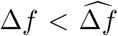), in that case the inner and outer repeat loops stop and the current **Z** (not **Z***_new_*) is outputted.

∇*f* (**Z**) is given by:

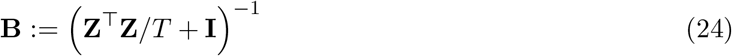

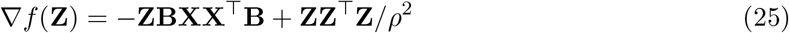

Finally, the expression for **Y** is (**Supplementary Information**, equation (S48)):

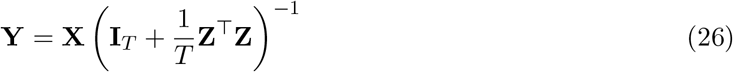

### Numerical simulation of the NNC offline

For the NNC, we do not have the analytical expressions of **Y** and **Z**. To minimize the objective function, we perform alternating gradient descent/ascent steps on **Y** and **Z**, respectively. We start from the expanded expression of the objective function (18) (with *γ* = 1):

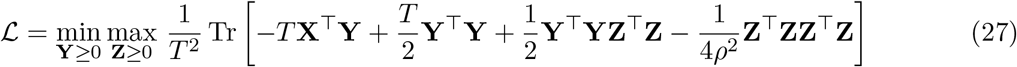

#### Algorithm 2 Finding the minimum in Y and maximum in Z of 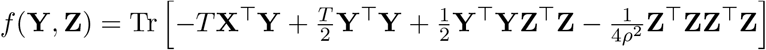

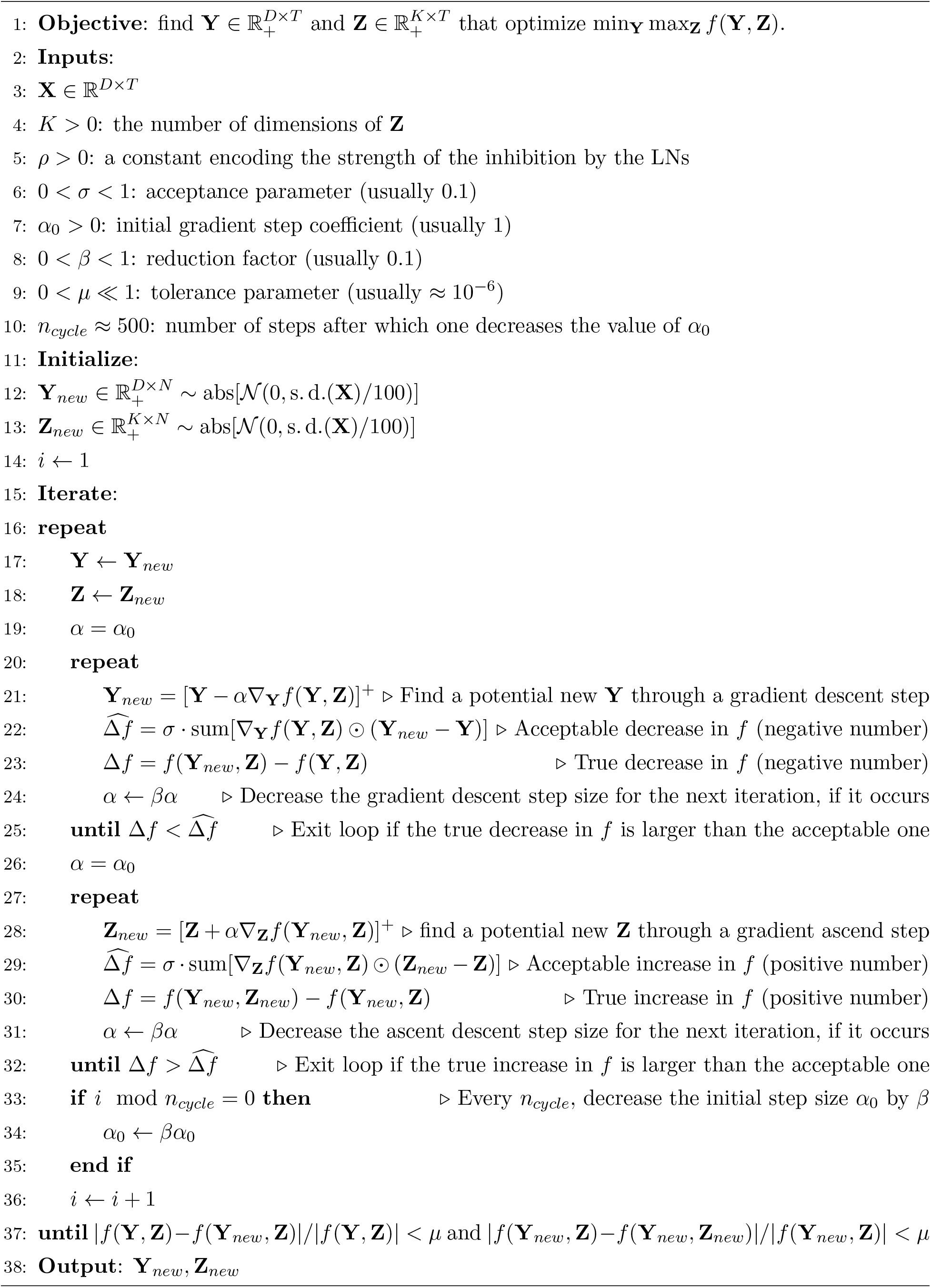

In the case of the LC, the same algorithm holds, with all the rectifications [.]^+^ removed from the algorithm and the “abs” removed from the initiation. If in either of the inner repeat loops, no α is small enough to make a decrease/increase in *f* (i.e., satisfy the condition 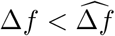 or 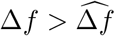), the iterations stop and the current **Y** and **Z** are the output of the algorithm.

The gradients of *f*(**Y, Z**) are:

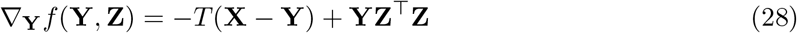

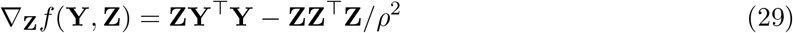

### Numerical simulation of the circuits online

For **Fig. S11**, we simulated the circuit dynamics for a given **W, M**, and **X**. For that purpose, to find 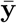 and 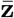, we performed gradient descent steps based on the discretized equations (19) for the LC or equation (20) for the NNC.

## Data and code availability

All data in this study is published in Berck et al., 2016; Si et al., 2019 and is accessible online: https://github.com/samuellab/Larval-ORN, https://doi.org/10.7554/eLife.14859.019, https://doi.org/10.7554/eLife.14859.020.

All the code used in this study is available here: https://github.com/chapochn/ORN-LNcircuit

## Acknowledgments

We thank Aravinthan D.T. Samuel, Jacob Baron, Guangwei Si, Thomas Frank, Victor Minden, Anirvan Sengupta, Eftychios A. Pnevmatikakis, Shiva GhaaniFarashahi, and the Neuroscience Group at the Flatiron Institute for discussions and/or comments on the manuscript.

## Author contributions

All authors designed the study. C.P. and D.B.C. formulated the objective function. N.M.C. and C.P. performed theoretical derivations. N.M.C. wrote the computer code, analyzed the data, performed numerical simulations, and prepared the original draft. All authors reviewed and edited the manuscript.

## Supplementary Figures

**Fig. S1.**
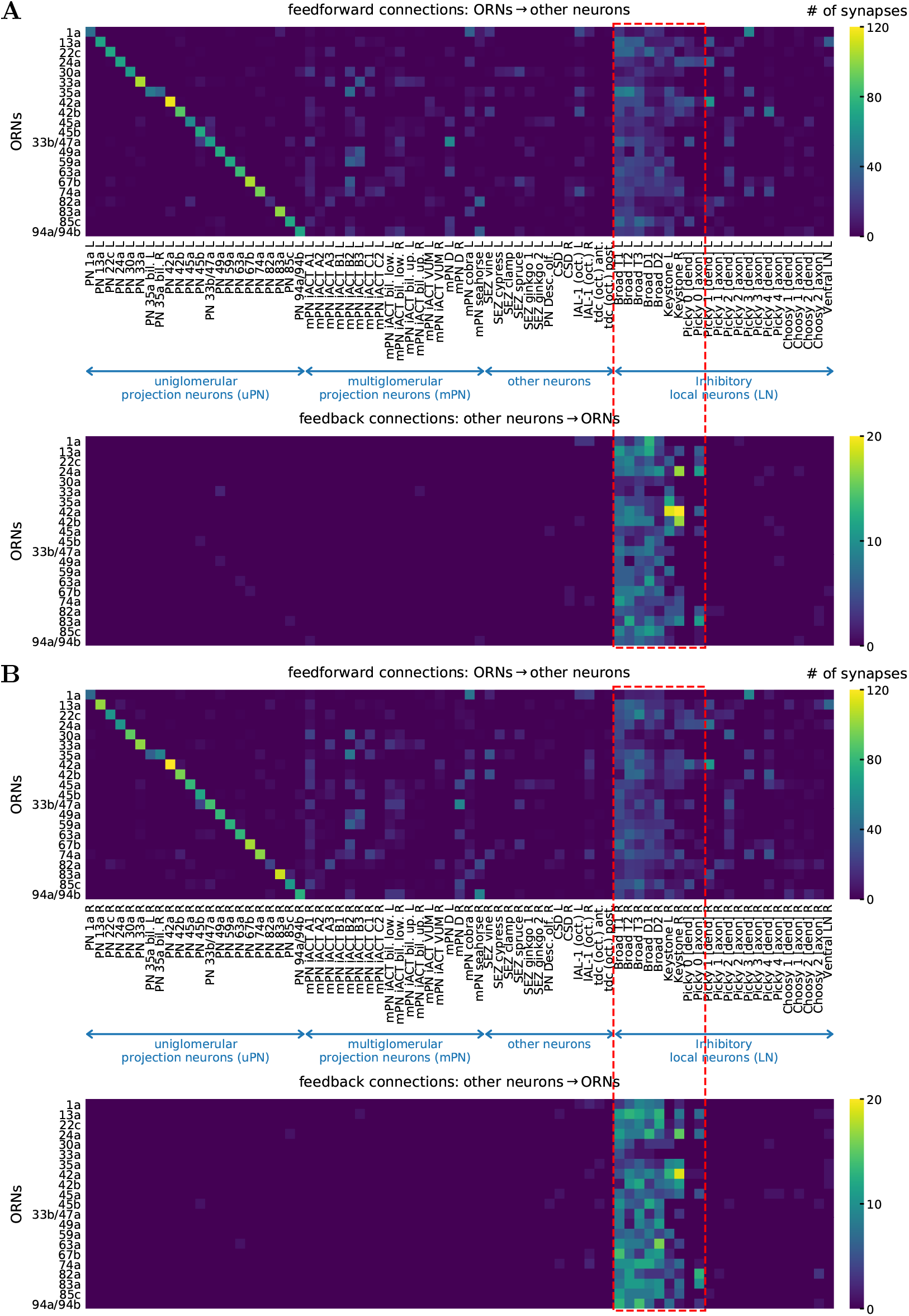
Full ORN connectivity and circuit selection. **A** Heat map of the ORNs ↔ LN feedforward and feedback connections on the left side of the *Drosophila* larva. We focus on the neurons, that synapse bidirectionally with ORNs (inside the red dashed rectangle): Broad Trios, Broad Duets, Keystones, and Picky 0. These neurons are all LNs. **B** Same as **(A**) for the right side.

**Fig. S2.**
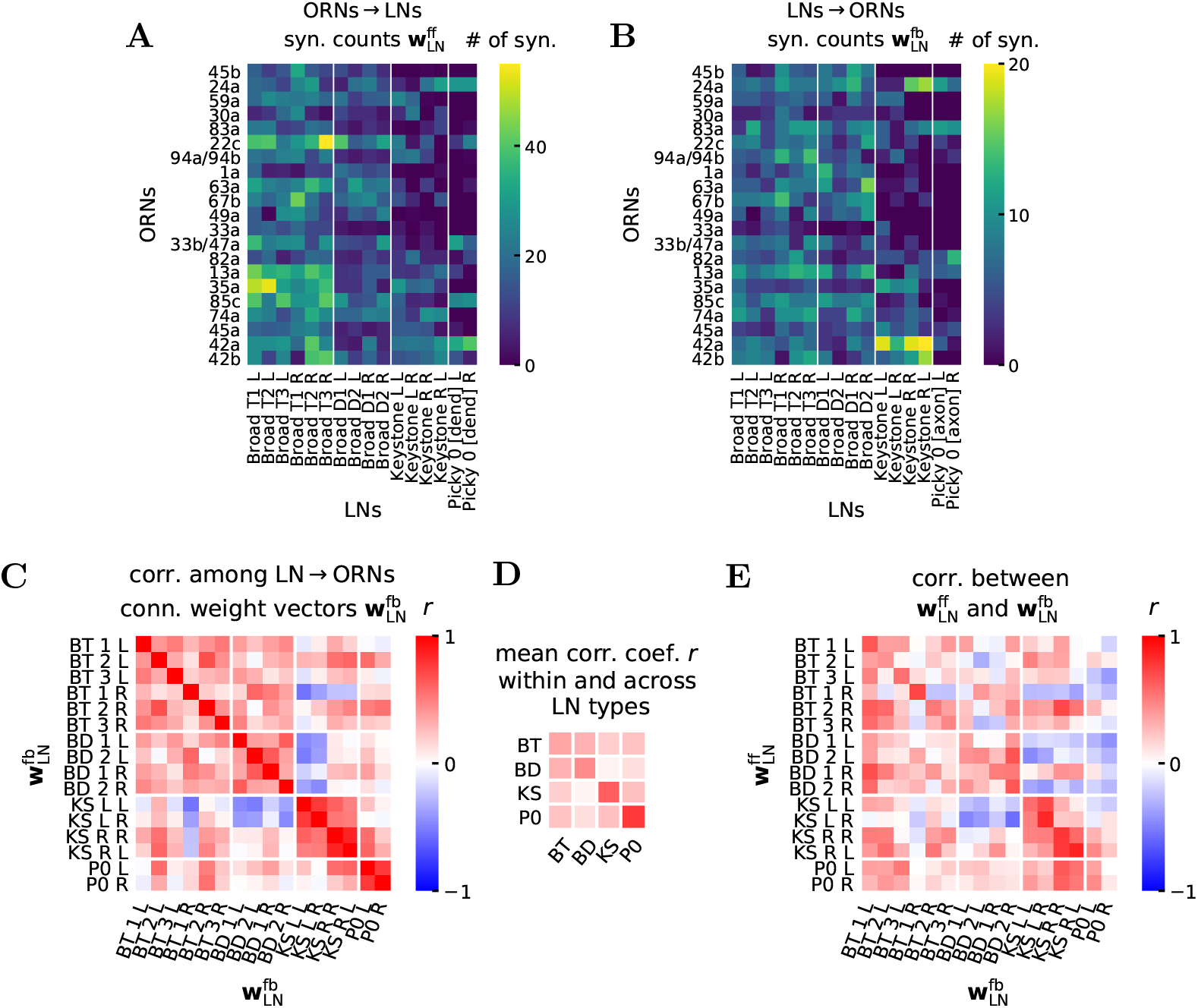
ORN-LN connectivity, comparison feedforward with feedback. **A** ORNs → LNs feedforward connections weights 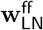 on both left and right sides of the antennal lobe with the chosen LNs, ordered by LN class. The vectors 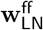 correspond to the columns of the depicted matrix. **B** LN → ORNs feedback connections weights 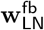 on both left and right sides of the antennal lobe with the chosen LNs, ordered by LN class. The vectors 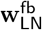 correspond to the columns of the depicted matrix. **C** Correlation coefficients between feedback LN → ORNs connection weight vectors 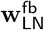. **D** Average rectified correlation coefficient 〈*r*_+_〉 (*r*_+_:= max[0, *r*]) between LN types calculated by averaging the rectified values from (**C**) in each rectangle with white border, excluding the diagonal entries of the full matrix. The average correlation coefficient within a class is larger than the correlation coefficient across classes. **E** Correlation coefficients between feedforward ORNs → LN 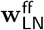 and feedback LN → ORNs 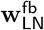 connection weight vectors. The Picky 0 LN is the only LN that has a separation between axonal and dendritic terminals. For the feedforward ORNs → LN connections, we only include in the connection weight vector the synapses onto the Picky 0 dendrite, and for the LN → ORNs connection, we only count the synapses from the Picky 0 axon.

**Fig. S3.**
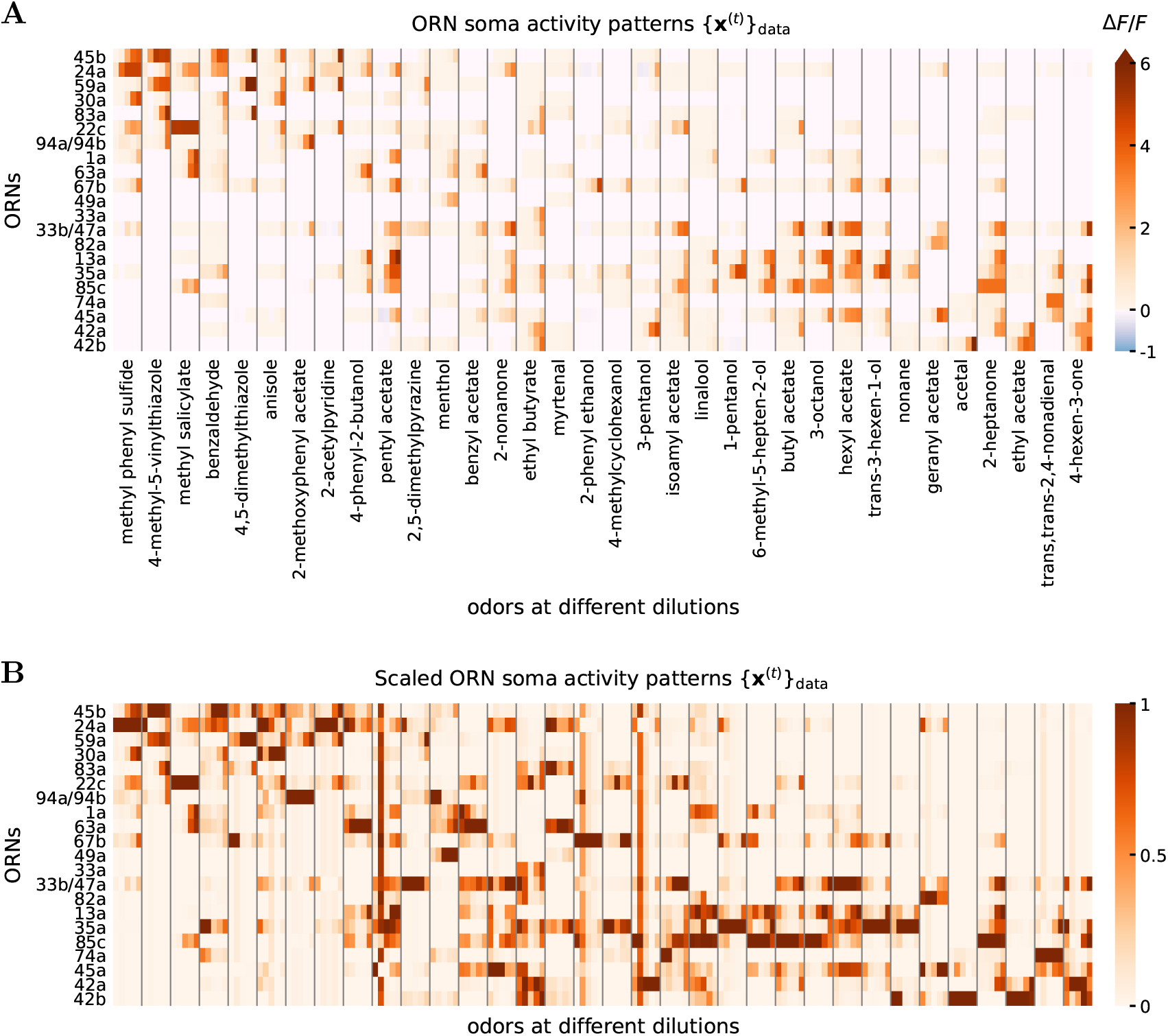
ORN soma activity from Si et al., 2019. **A** ORN soma activity patterns {**x**^(*t*)^}_data_ in response to 34 odors at 5 dilutions acquired through Ca^2+^ imaging. Different odors are separated by vertical gray lines. For each odor, there are 5 columns corresponding to 5 dilutions: 10^−8^,…, 10^−4^. The odors and ORNs are ordered by the value of the second singular vectors of the left and right SVD matrices of this activity data, after centering and normalizing. This data is obtained by averaging the maximum responses of several trials to the same odor and dilution (as in Si et al., 2019). **B** Same as (**A**), with each **x**^(*t*)^ scaled between 0 and 1 to better portray the patterns.

**Fig. S4.**
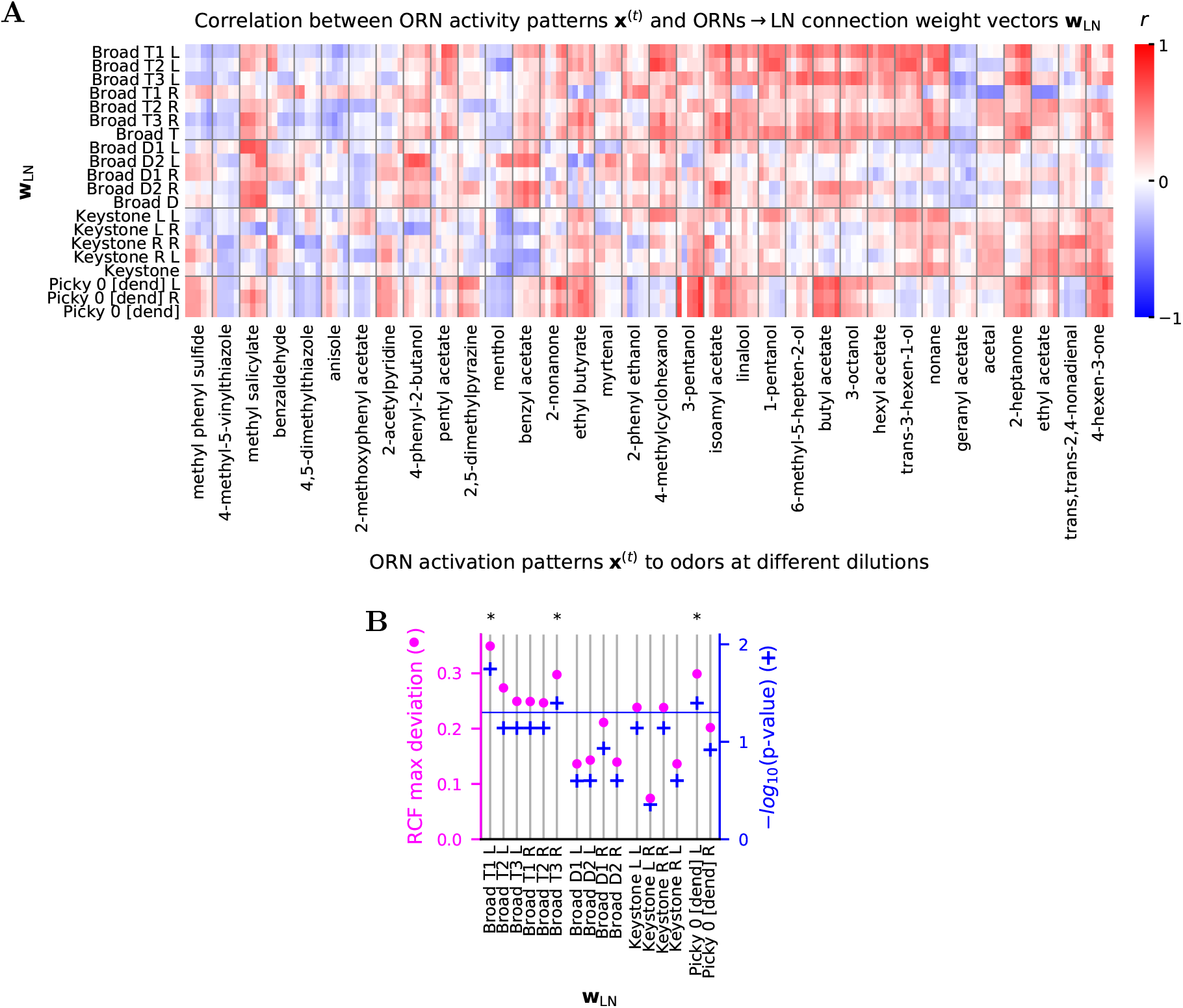
Alignment of activity patterns x^(*t*)^ in ORNs and ORNs → LN connectivity weight vectors w_LN_. **A** Sa me as **Fig. 2E**, for all the **w**_LN_ and with all the odors labeled. Same odor order. **B** Sa me as **Fig. 2I**, for all **w**_LN_.

**Fig. S5.**
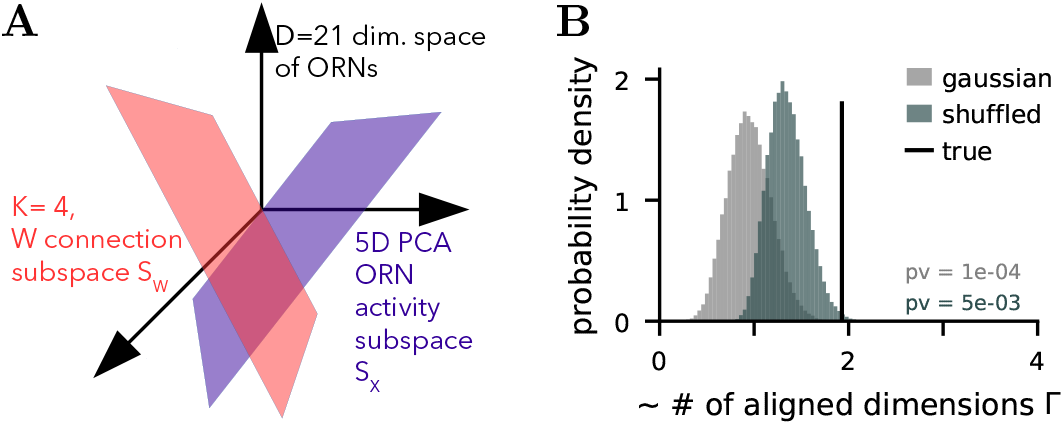
Activity and connectivity subspace alignment. **A** Scheme representing the comparison of the 4-dimensional connectivity (*S_W_*) and 5-dimensional activity (*S_X_*) subspaces in 21 dimensions (*D* = 21, dimensionality of the ORN space). **B** Number of aligned dimensions Γ between the 2 subspaces of (**A**) in the data (true, Γ = 1.9), from randomly shuffling the connectivity vector entries (shuffled, mean Γ = 1.3) and from random normal vectors (Gaussian, mean Γ = 1). pv: one-sided p-value.

**Fig. S6.**
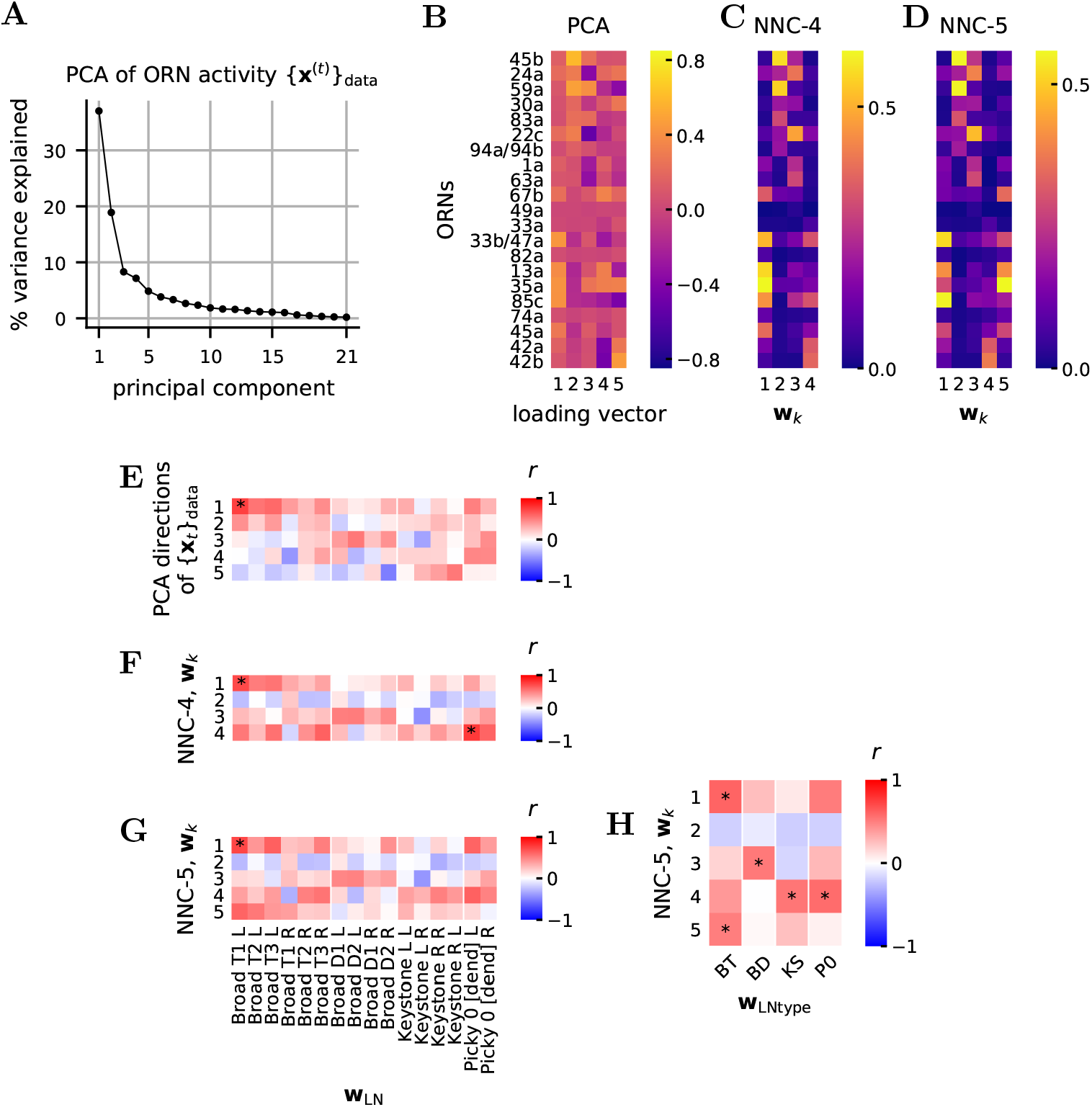
Activity and connectivity. **A** Percentage of the variance of the ORN activity patters {**x**^(*t*)^}_data_ explained by the uncentered PCA. The top 4 and 5 PCA directions explain 71% and 76% of the variance, respectively. **B** First 5 PCA loading vectors of {**x**^(*t*)^}_data_. **C-D w**k from NNC with *K* = 4,5 and *ρ* =1, ordered to resemble the PCA ordering. **E** Same as **Fig. 3C** with all **w**_LN_. **F** Same as (**E**), with **w***_k_* from NNC-4 instead of PCA loading vectors. **G** Same as (**F**), for NNC-5. The small number of significant points in (**E-G**) results from the higher number of hypothesis tests, which decreases the adjusted p-values in the FDR multi-hypothesis testing framework. **H** Same as **Fig. 4A**, for NNC-5.

**Fig. S7.**
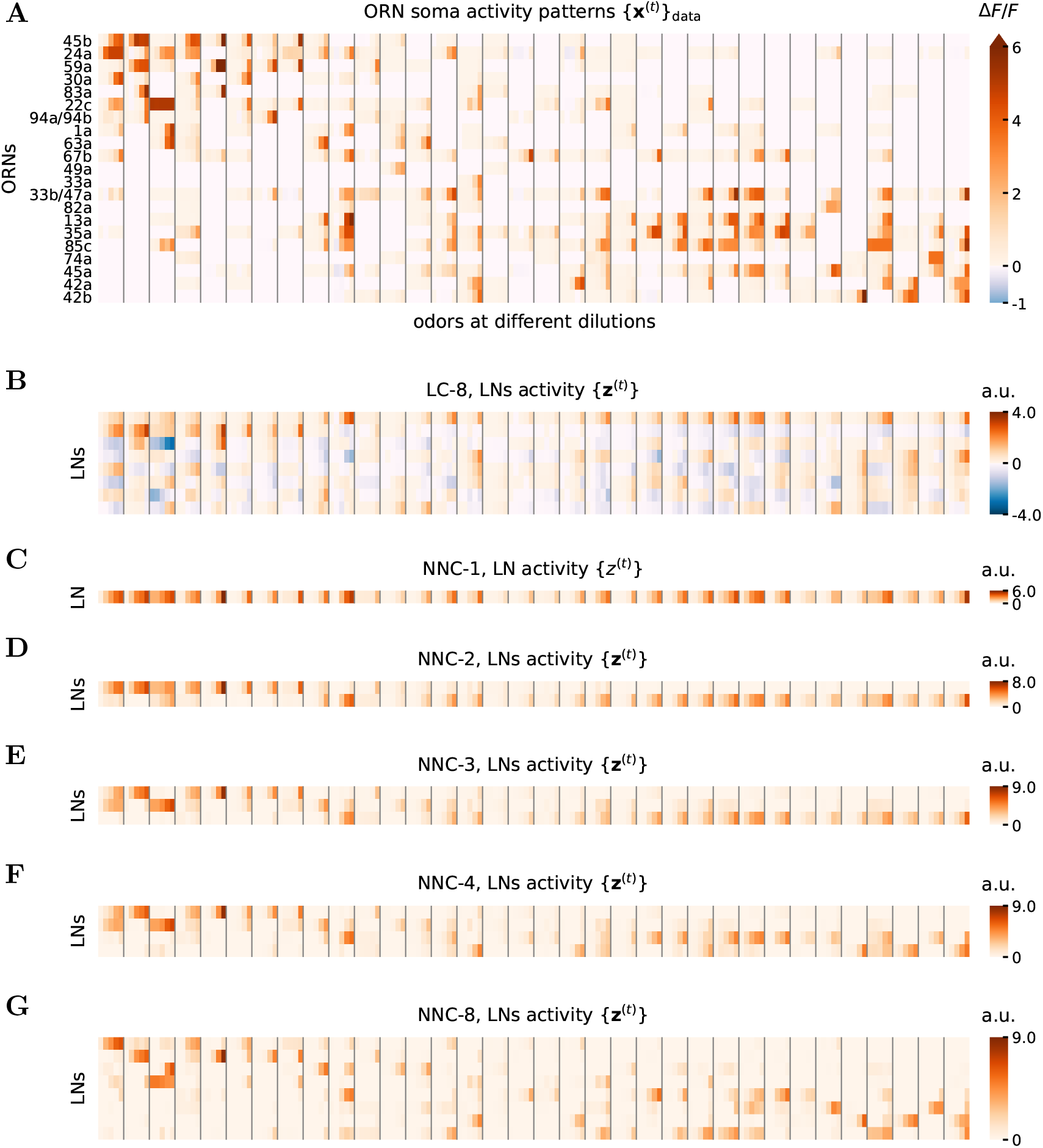
Activity of LNs {**z**^(*t*)^} in the NNC and LC. **A** ORN soma activity patterns {**x**^(*t*)^}_data_ as in **Fig. S3A**. **B** Activity in the LNs for the LC-8. Stimuli are aligned to the panel above. As mentioned in the text, {**z**^(*t*)^} is undetermined up to an orthogonal matrix **U***_Z_*. Here we set **U***_Z_* = **I***_K_*, i.e., identity matrix. For LC-*K*, the response in LNs correspond to the first *K* row of this matrix, multiplied by any *K* × *K* orthogonal matrix on the left. Thus, the matrix depicted in this plot shows the potential activity in LNs for any LC-K with *K* ≤ 8. **C** {*z_t_*} for the NNC-1. The activity of the LN approximately follows the total activity. **D** {**z**^(*t*)^} for the NNC-2. One can see that the 2 LNs roughly clusters the sets of odors into those activating the top ORNs and those activating the lower ORNs. **E-G** {**z**^(*t*)^} for the NNC with *K* = 3,4,8. One observes a more sophisticated clustering of the data. As more LNs are added, LN activity increases in sparsity. The activity in the LNs for the NNC is more sparse than for the LC.

**Fig. S8.**
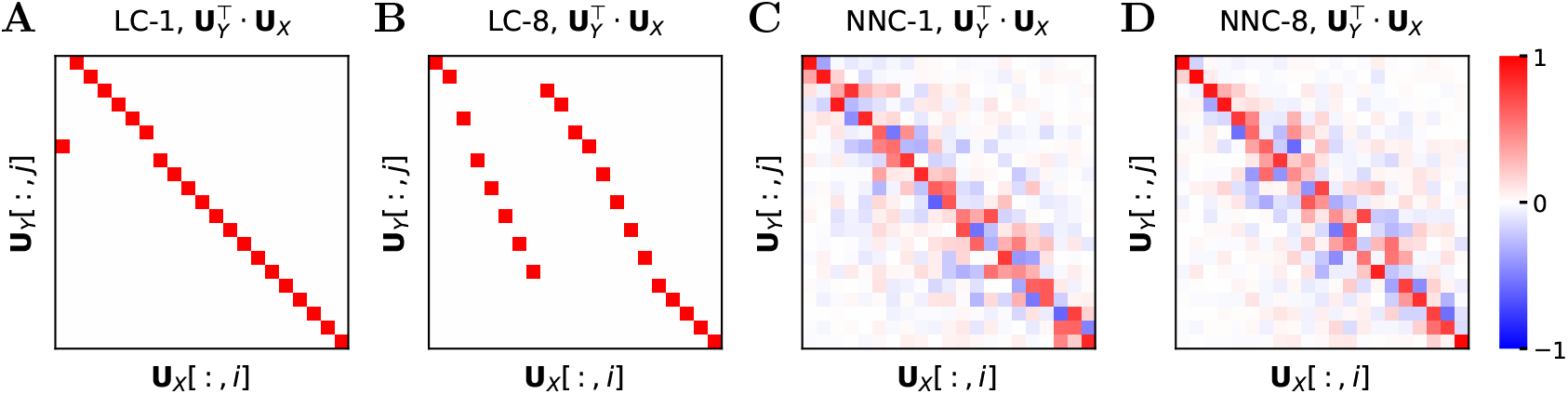
Input vs output principal directions in LC and NNC. **A-D** Scalar product between principal directions of uncentered {**x**^(*t*)^}_data_ and {**y**^(*t*)^} for the LC and NNC for *K* = 1,8. For the LC the identity of the principal directions in conserved, only their order change. For the NNC, the principal directions are slightly mixed, but conserve the approximate ordering.

**Fig. S9.**
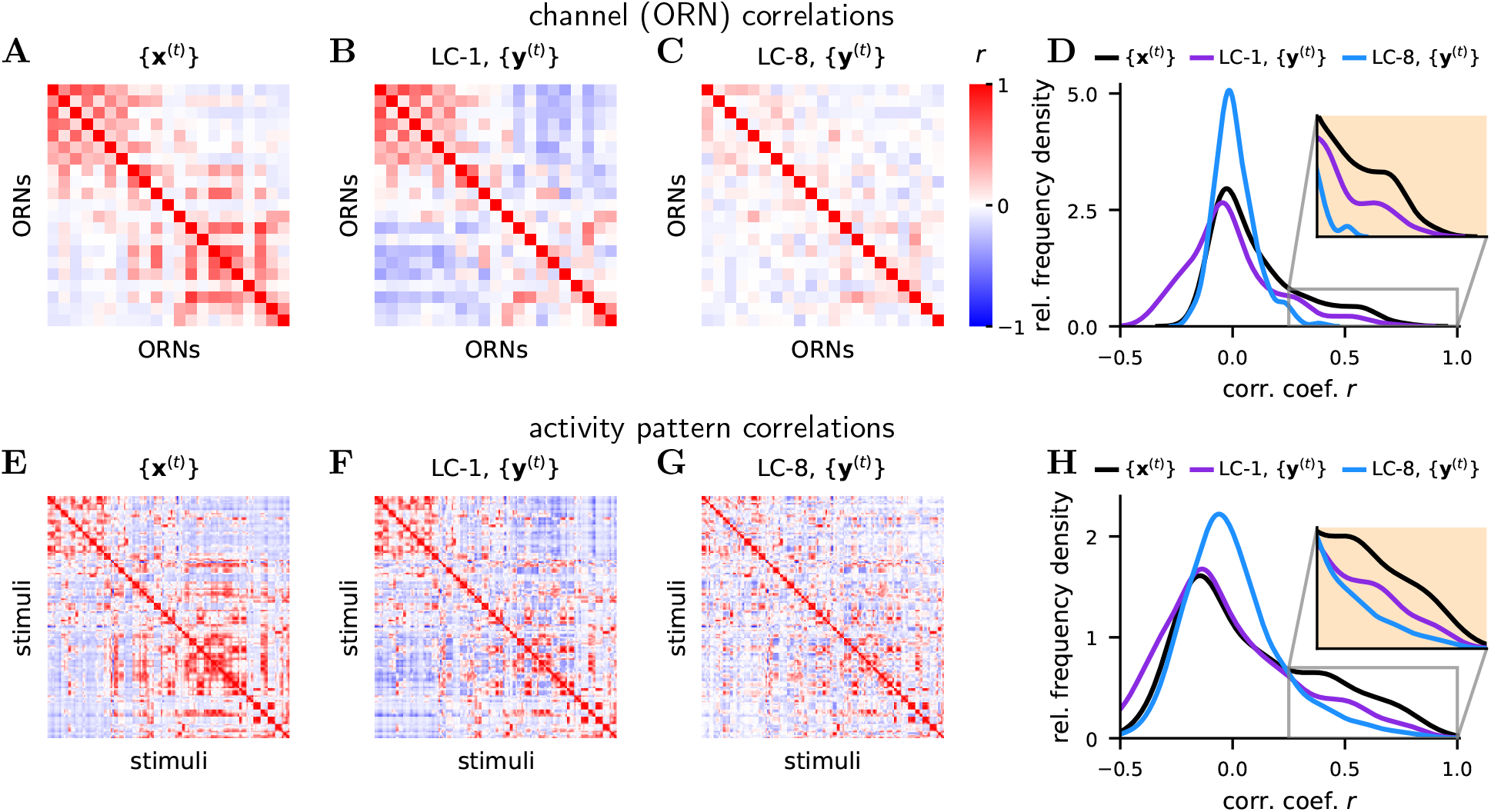
Decorrelation in the LC. **A-H** Same as **Fig. 7J-O** for the LC.

**Fig. S10.**
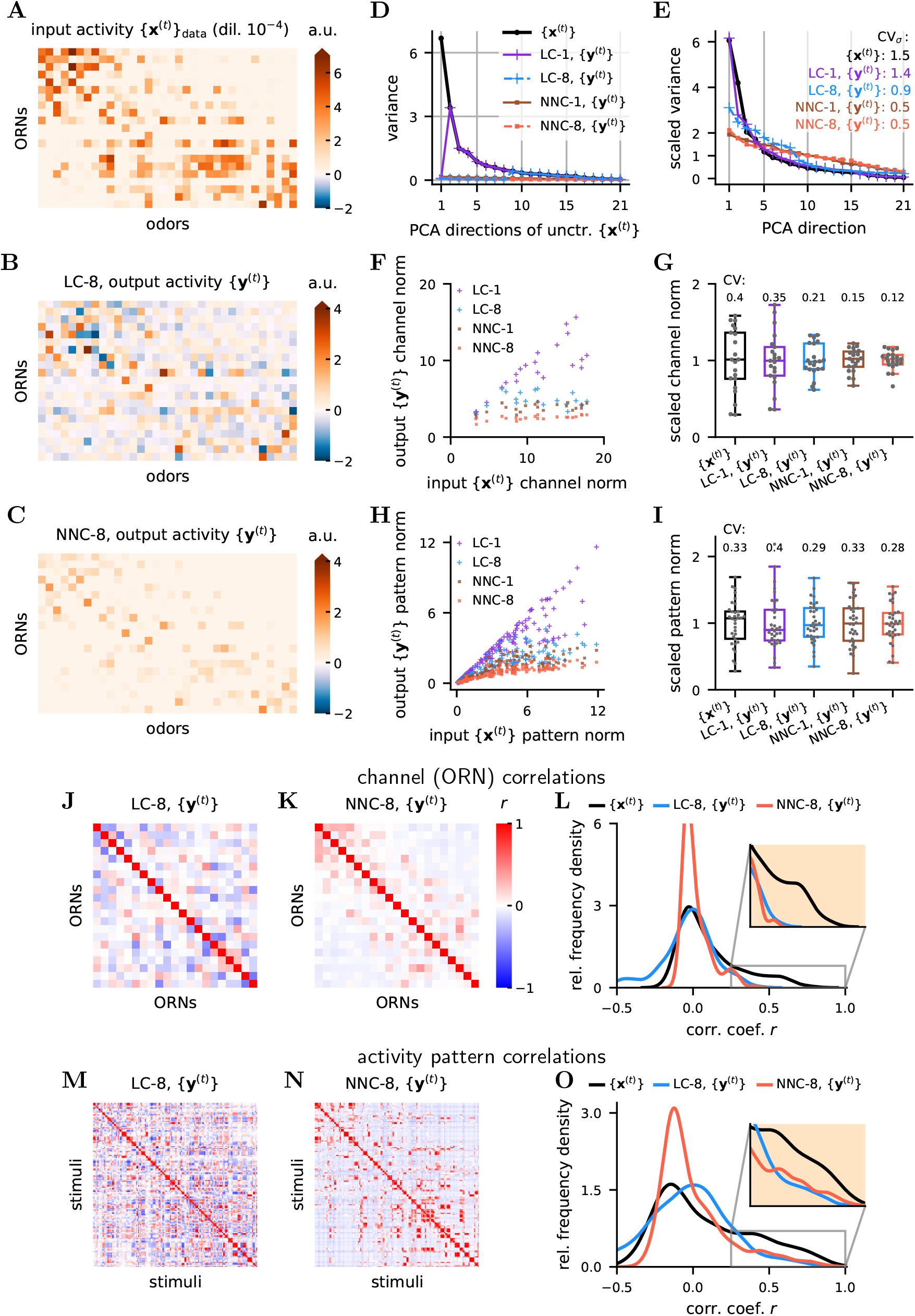
Input transformation by LC and NNC with *ρ* = 10. Same as **Fig. 7** for *ρ* = 10. Note the even stronger dampening, flattening, and decorrelation.

**Fig. S11.**
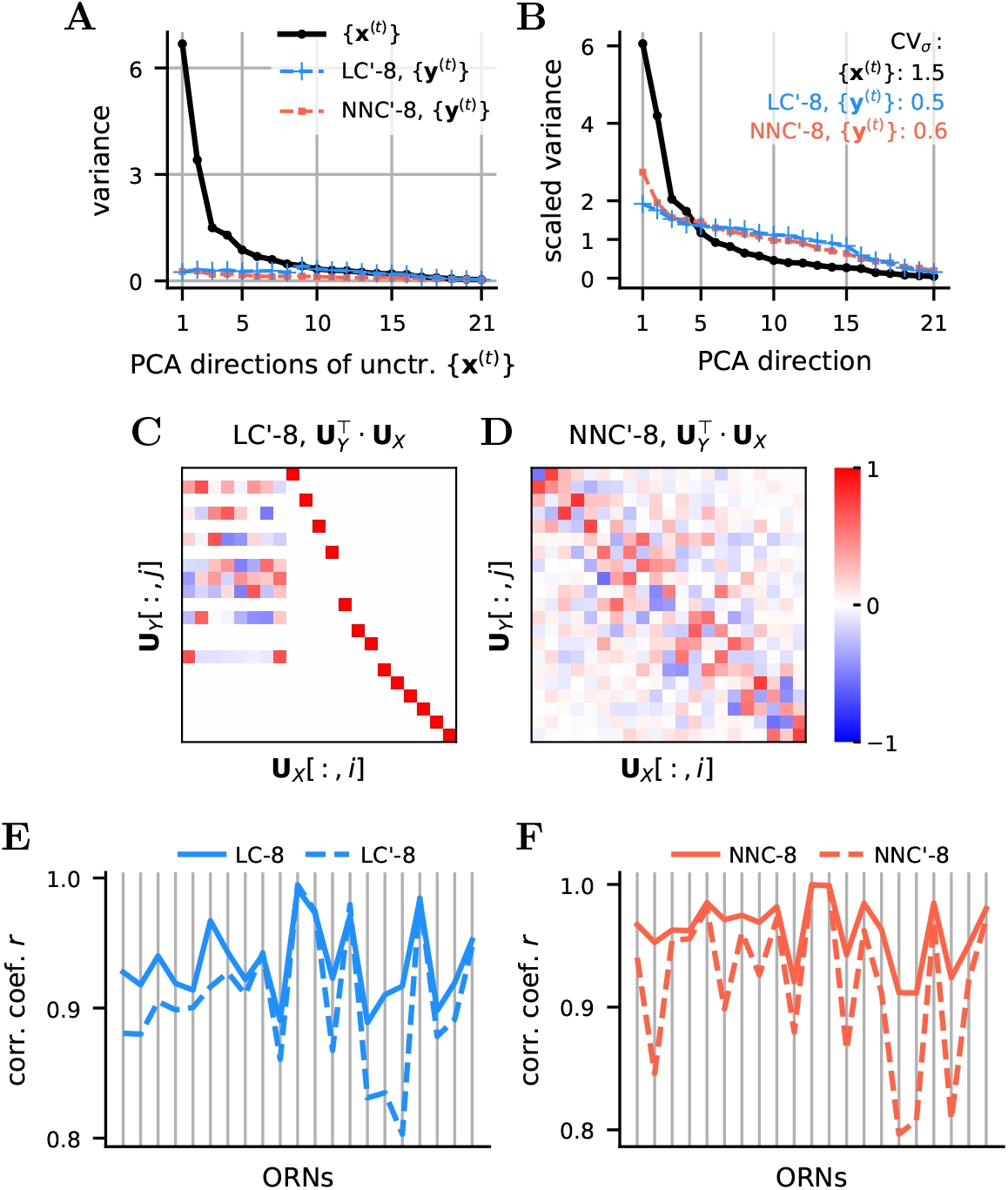
Effect of removing off-diagonal entries in M for LC and NNC. **A-B** Same as **Fig. 7D,E** for the trained LC and NNC on {**x**^(*t*)^}_data_, where the off-diagonal values of **M** are set to 0 (LC’ and NNC’). Note that the values in LC’ in (**A**) do not monotonically decrease as in LC. **C-D** Same as **Fig. S8** for LC’-8 and NNC’-8. Note the increased mixture between the principal directions of {**x**^(*t*)^}_data_ and {**y**^(*t*)^}. **E** Correlation between the input {**x**^(*t*)^}_data_ and output {**y**^(*t*)^} for each channel (i.e., ORN) for LC-8 and LC’-8. Note that in the LC-8, the output of each channel is more strongly correlated to its own input for the LC-8 than for the LC’-8. **F** Same as (**E**) for NNC-8 and NNC’-8.

## Supplementary Information

In this supplement, we prove statements made in the results and methods sections:

Section 1: we describe the objective function from equation (18), show the equivalence of scaling **X** and *ρ* (section 1.1) and show the resemblance of this circuit’s objective function with a whitening objective function (section 1.2).
Section 2: we show that the objective function (18), when optimized online with or without non-negativity constraints, lead to the circuit dynamics (19) or (20), respectively, and to Hebbian learning rules (21). We then show the steady state solution to which the circuit dynamics equations (19) converge and show that the steady state is stable (section 2.2).
Section 3: we show that a general system of differential equations describing the circuit contains two effective parameters and can be reduced to the form found in the main text in equation (19).
Section 4: we analyze computation in LC and prove equations (4), (5), (6a), and (6b) in the main text from the main text. These results are proven in two ways (sections 4.1 and 4.2). Section 4.3 discusses limiting cases of the computation for small and large values of ρ, and show the relation of NNC to SNMF (symmetric non-negative matrix factorization).
Section 5: we prove the relationship between **W** and **M**, equation (2) in the main text.
Section 6: we prove the relationship between **W** and **X**, equation (1) in the main text.
Section 7: we prove that the CV of singular values in **Y** is smaller than in **X** for the LC when *K* = *D*.

### 1 Objective function

We postulate the following minimax objective function:

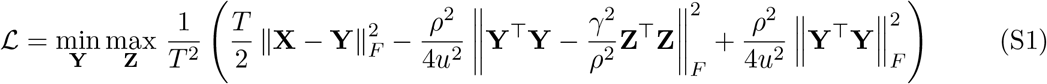

Which can be expanded thus:

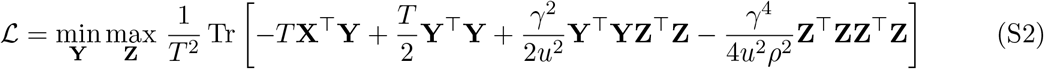

Where **X**, 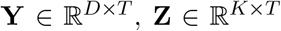 with *D* the number of ORNs (21 for this olfactory circuit), *K* the number of LNs, *T* the number of data (sample) points, *ρ* a positive unitless constant, *u* a unit with the physical dimension as **X, Y**, and **Z** (e.g., spikes o s^−1^) (dropped for simplicity in the main text). **X, Y** and **Z** represent the activity of ORN somas, ORN axons, and LNs, respectively. We can interpret **X** as all the discretized activity of ORNs up to a certain point in their lifetime.

Optimizing objective function (S2) leads to the linear circuit (LC) model. Adding the non-negativity constraints on **Y** and **Z** leads to the non-negative circuit (NNC) model.

#### 1.1 Equivalence of scaling X and *ρ*

Here, we show that scaling **X** is equivalent to scaling *ρ* in terms of circuit computation. It is easy to see that the transformation **X** → *a***X, Y** → *a***Y** and *ρ* → *ρ/a* (for *a* ≠ 0) leaves the objective function unaffected, i.e., this transformation is a symmetry of the optimization. Indeed:

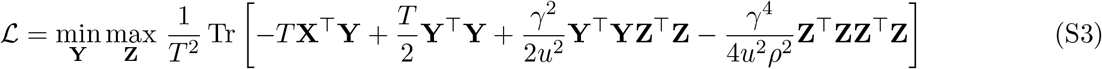

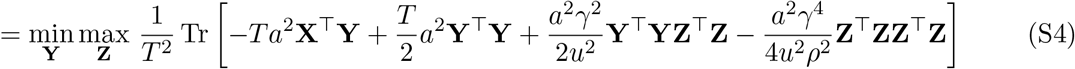

Let us explore the consequence of this symmetry. The output **Y** of the optimization is a function of **X** and *ρ*, thus we can define a function *f* such that: **Y** = *f*(**X**, *ρ*):

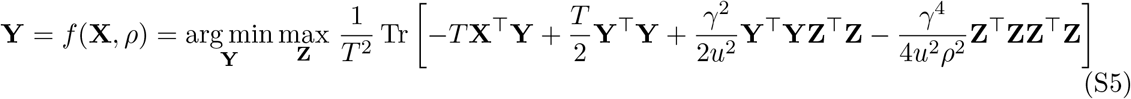

The symmetry implies:

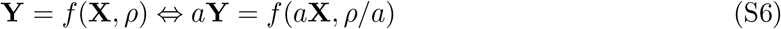

Thus:

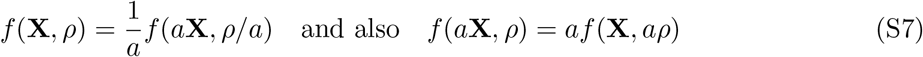

This means performing an optimization with an input *a***X**, is equivalent to doing the optimization with input **X** and parameter *aρ*, and finally multiplying the obtained **Y** by *a*.

It is worth noting though, that for a circuit with fixed **W** and **M**, scaling an input **x** by a factor *a*, simply scales the output **y** by the same factor a, since it is a linear transformation, at least for the circuit without the non-negative constraints.

#### 1.2 Limiting case and relation to whitening

For the case when *D* = *K*, the optimum for **Z** is 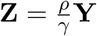 and thus the middle term of the objective function (S1) drops, with and without non-negativity constraints on **Y** and **Z**. The objective function becomes:

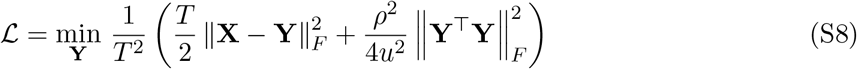

This objective function closely resembles the whitening objective function:

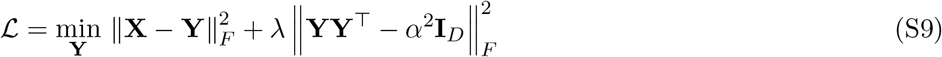

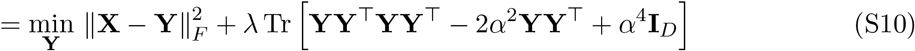

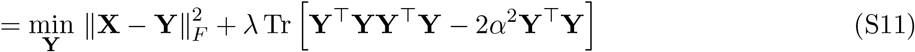

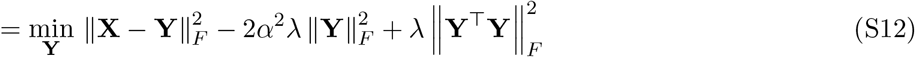

For a fixed *α*, increasing λ will eventually lead to perfect whitening. The singular values of **Y** will then all be equal to *α*, and the left and right singular vectors will be the same as those of **X**.

### 2 Online solution

We show that the online algorithm to optimize the objective function (S2) can be mapped onto the architecture and neural dynamics of the olfactory neural circuit (**Fig. 1A**) with Hebbian plasticity. To find the online solution, we first introduce the unitless variables 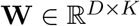 and 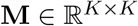:

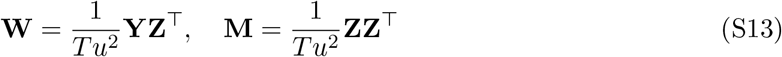

and perform the Hubbard-Stratonovich transform of (S2):

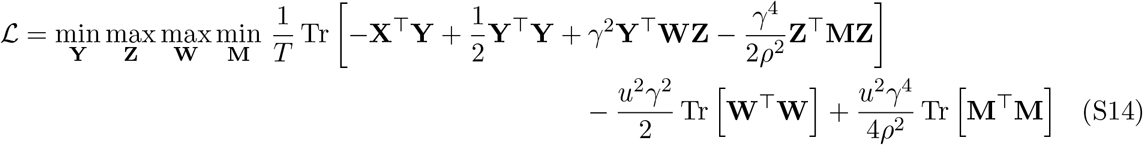

We then rewrite (S14) in vector notation, with each sample point written out separately, and invert the order of min max (Pehlevan et al., 2018):

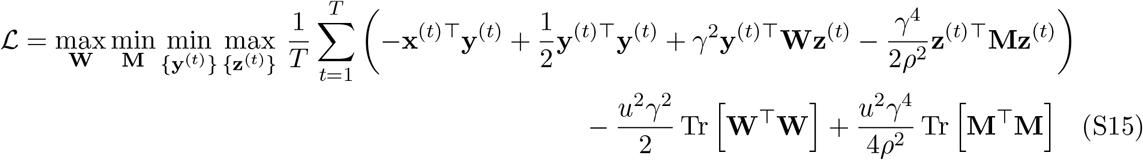

Next we perform the optimization for each variable separately: **y**^(*t*)^, **z**^(*t*)^, **W**, and **M**. We consider the following case, which corresponds to the “online setting” for this objective function and alternate the optimization in {**y**^(*t*)^, **z**^(*t*)^} and in {**W, M**}: as a new sample (i.e., stimulus, input) **x**^(*t*)^ arrives, we find the values of **z**^(*t*)^ and **y**^(*t*)^ with the current values **W**^(*t*)^ and **M**^(*t*)^ and update **W**^(*t*)^ and **M**^(*t*)^ to **W**^(*t*+1)^ and **M**^(*t*+1)^ before the arrival of the next sample **x**^(*t*+1)^. Biologically, this can be seen as first a convergence of neural spiking rates or neural electrical potential encoded through the variables **y**^(*t*)^ and **z**^(*t*)^, and second a synaptic weight update based on those steady state activity values. At a given sample index *t*, the minimum in **y**^(*t*)^ and the maximum in **z**^(*t*)^ can be found by taking a derivative of (S15) with respect to **y**^(*t*)^ and **z**^(*t*)^, respectively:

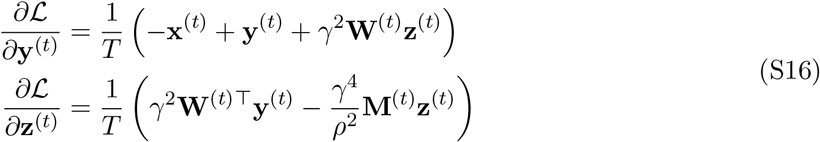

The minimum in **y**^(*t*)^ and the maximum in **z**^(*t*)^ can be reached by a gradient descent and ascent, respectively. We can thus write a system of differential equations whose steady state correspond to the optimum:

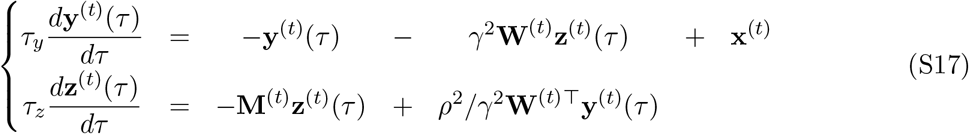

Where *τ* is the local time evolution variable. We rearranged the parameters so that the equation form is the same as in equations (19), which does not change the final steady state of the equations. Thus, we obtained equations to find the optima 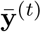 and 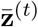 of the objective function. As explained in the main text, these question can directly be mapped onto the dynamics of the ORN-LN neural circuit.

Next, we derived the updates for the variables **W** and **M**. By construction, the offline solution for **W** and **M** is given by (S13). Online - we compute a new **W**^(*t*)^ and **M**^(*t*)^ after each sample **x**^(*t*)^ is presented and 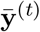 and 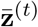 are found. The gradient descent (respectively ascent) steps on these variables give the following updates (e.g., Pehlevan et al., 2018):

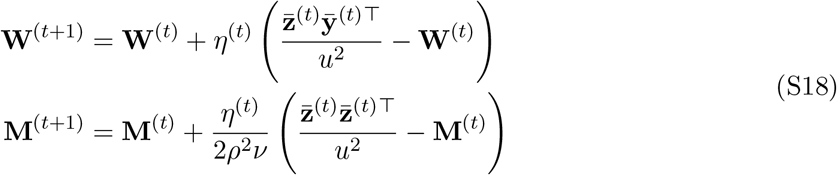

where and *η*^(*t*)^ and *ν* are parameters of the gradient descent/ascent, and where 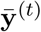 and 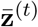 are the steady states solutions of equations (S17) for given **W**^(*t*)^ and **M**^(*t*)^. This indeed corresponds to a local Hebbian synaptic update rules. Choosing *η*^(*t*)^ and *ν* appropriately will lead to equation (21) from the main text.

#### 2.1 Circuit equations for the NNC

In the case of the NNC, where we have objective function (18) instead of (S2), we get equation (S15) with non-negativity constraints:

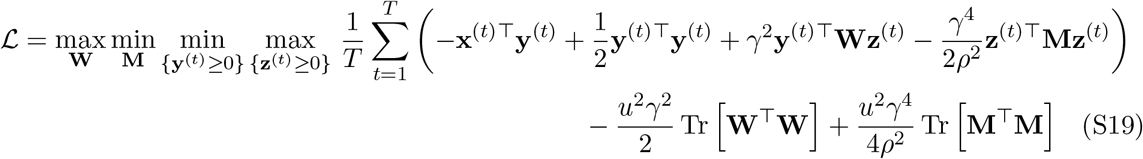

Here too, we perform the optimization for each variable separately: **y**^(*t*)^, **z**^(*t*)^, **W**, and **M**. However, because of the non-negativity constraints, the optima for **y**^(*t*)^ and **z**^(*t*)^ are not to be found at where the derivatives (S16) are zeros. We can, however, reach the optima by a projected gradient descent:

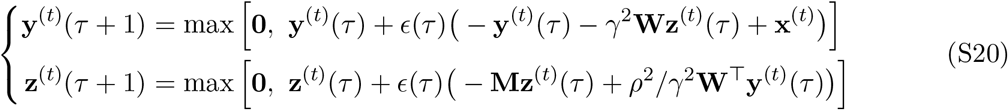

where the max is performed component-wise. For the NNC, the updates on **W**^(*t*)^ and **M**^(*t*)^ (equations (S18)) remain the same as for the LC.

#### 2.2 Steady state solution of the circuit dynamical equations for the LC and stability

We can directly find the steady state solution of the circuit dynamics equations (S17) of the LC by setting the derivatives to 0. For **M** invertible, the steady state is (after dropping the index (*t*) for simplicity of notation):

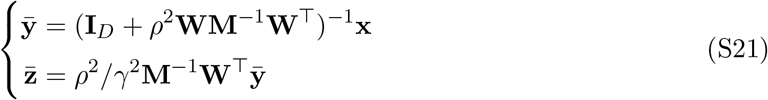

As mentioned above, the steady state for **y** does not depend on *γ*, whereas **z** does depend on *γ*. Note that the transformation from **x** to 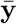 is symmetric: indeed, writing 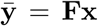, we have that **F** = **F**^⊤^. This means that the transformation is diagonalizable. It will be shown below that this basis in which the transformation is diagonal corresponds to the PCA basis of **X**.

Here we show that the fix point of equations (S17) is stable if **W** is maximum rank and **M** positive definite. We first rewrite the dynamical system:

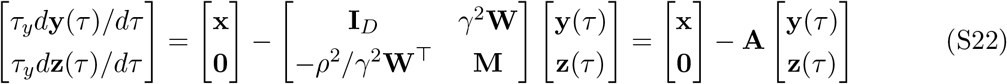

This system has a unique stable fix point if and only if **A** has only positive eigenvalues. To investigate under which conditions this is the case, we write the eigenvalue equations for **A**:

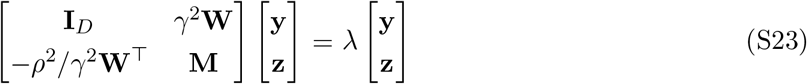

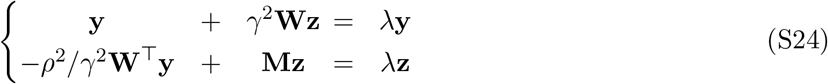

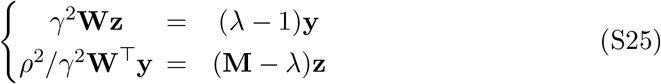

We consider the case when λ ≠ 1, as we are interested to see if λ could potentially be negative.

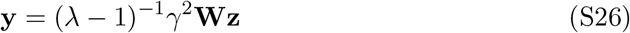

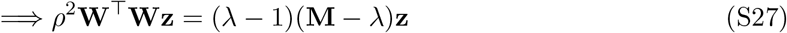

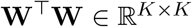 is a positive semi-definite matrix, it is positive definite if **W** is maximum rank (i.e., rank *K*). Assuming that **W** is full rank, the matrix **W**^⊤^**W** on the left-hand side of the equation has only positive eigenvalues. The above equation does not have any solution **z** ≠ **0** for λ < 0 if **M** is positive definite (which is true when constructed as the autocorrelator of **z**). Thus, **W** full rank and **M** positive definite are sufficient conditions for the dynamical system to always converges to a stable fix point.

### 3 Circuit dynamics equations contains two effective parameters

Here we show that, in its general form, the system of differential equation describing the olfactory circuit has just two effective parameters and can be reduced to equation (19) (or (20)) from the main text. Without lack of generality the system of differential equations yields:

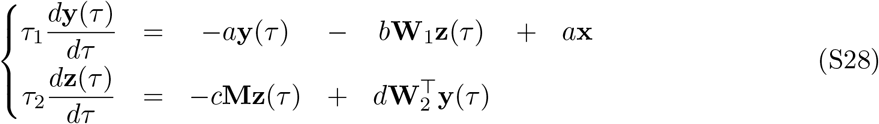

Where we imposed that **x** = **y** in the case of no LN activity (i.e., **z** = **0**), that *a* > 0, *b* > 0, *c* > 0, *d* > 0, and that all ORNs have similar response properties (i.e., same coefficient in front of each *x_i_* and *y_i_*). To extract the effective parameters, we compute the steady state solution of equations (S28) by setting the derivatives to zero. We find, for invertible **M**:

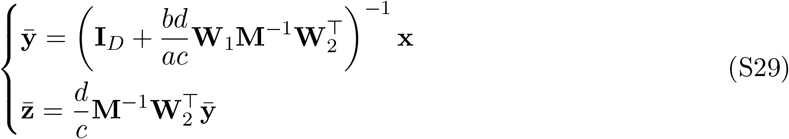

This shows that we only have two degrees of freedom: 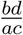 and 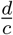. We define 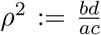 and 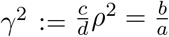. This gives us:

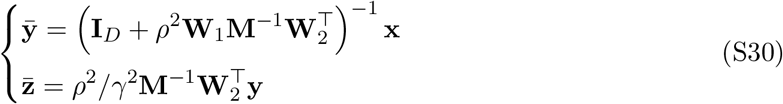

Now replacing these definitions into the original equations (S28) we get:

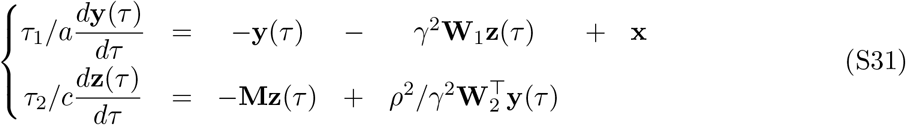

By setting *τ_y_*:= *τ*_1_/*α, τ_z_*:= *τ*_2_/*c* we obtain equation (19) from the main text (when **W**_1_ = **W**_2_):

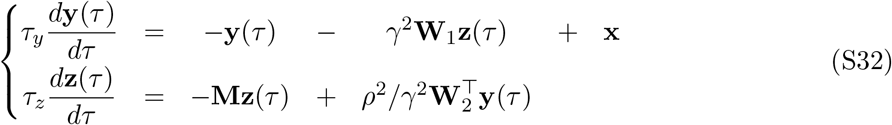

Thus, scaling **x, W**_1_, **W**_2_ and **M** is equivalent to controlling just two effective parameter *γ* and *ρ*. Scaling *τ_y_* and *τ_z_* does not influence the steady state solutions.

Increasing *ρ* increases the weight of feedforward connection, making the LN activity and the feedback inhibition stronger. Increasing *γ* simultaneously increases the feedback connection strength and decreases the feedforward connection strength. Changing *γ* influences the steady state solution of **z** but not **y**. Thus, a manifold of circuits lead to the same steady state output **y**. In addition, the same differential equations can be implemented by different circuits. For example, multiplying a differential equation by a parameter does not alter the final steady state, but gives yet another implementation to the circuit as a scaling of the synaptic weights and of the time constant.

### 4 Circuit computation

To understand the computation performed by the olfactory circuit, we analyzed the optimization done by the objective function (S2), which corresponds to the linear circuit (LC). We use the singular value decomposition (SVD) for **X, Y**, and 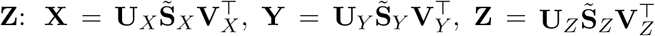, with the following convention: 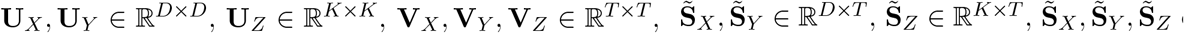 only have values on the diagonal. We call 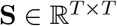 the diagonal square matrix corresponding to the rectangular matrix 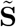, with padded zeros. Only the first *D* columns in **V***_X_* and **V***_Y_* and the first *K* in **V***_Z_* contain relevant information about **X, Y**, and **Z**, respectively. The left singular vectors **U***_X_*, **U***_Y_*, and **U***_Z_* are also the principal directions of the uncentered PCA of **X, Y**, and **Z**, respectively. Whereas the values on the diagonal of 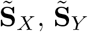, and 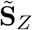 are the square root of the variances of the corresponding PCA directions.

In the following, using two approaches, we prove that:

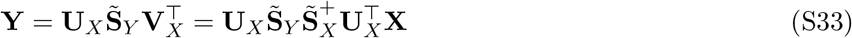

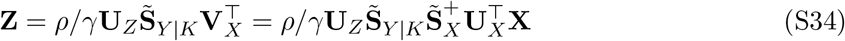

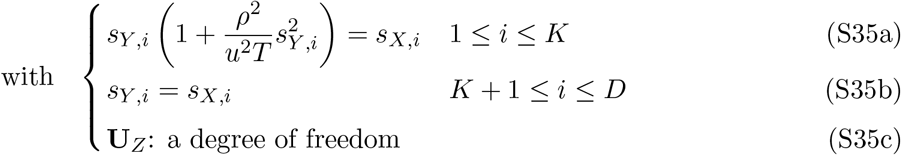

where **A**^+^ the Moore-Penrose pseudo-inverse of **A** and 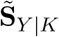 is the matrix with the first *K* columns of 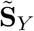. This proves the relations (4), (5), (6a), (6b) in the main text.

In other words, writing **Y** = **FX**, we have that 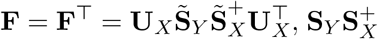 being a diagonal matrix. This signifies that the linear transformation **F** does not perform any rotation of the input.

This explicit expressions for *s_Y_* and *s_Z_* are:

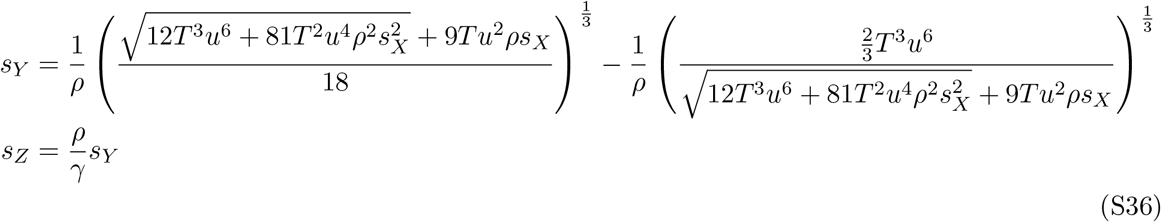

The behavior of *s_Y_* is such:

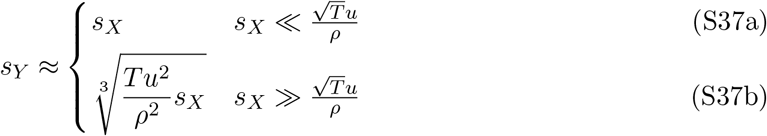

Note that because **Z** only appears as **Z**^⊤^**Z** in the objective function (S2), **U***_Z_* is a degree of freedom of the optimization. Thus, for {**Y, Z, W, M**} a solution of the optimization, {**Y, QZ, WQ**^⊤^, **QMQ**^⊤^} is a solution as well, where 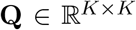 is an orthogonal matrix. Consequently, there is a manifold of **W, M**, and **Z** that satisfies the optimization for the LC.

#### 4.1 Approach 1

In this approach, we first perform the minimization in **Z**. Based on the similarity matching objective function results (Pehlevan et al., 2018), we know in the linear case that the right singular vectors of **Y** and **Z** are equal, and thus **V***_Y_* = **V***_Z_*. We also know that the top *K* singular values of **Y** and *γ/ρ***Z** are equal (**Z** is K-dimensional, thus all other singular values of **Z** are 0), and thus *s_Z,i_* = *ρ/γs_Y,i_*. The similarity matching term becomes:

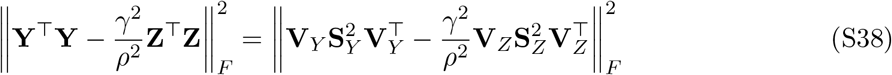

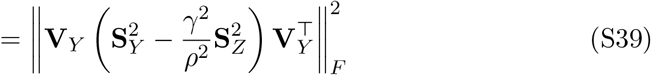

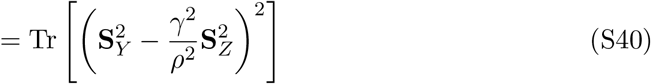

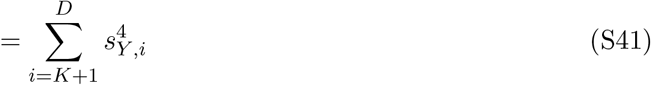

And thus **U***_Z_* does not appear in the optimization and is a free parameter. Also 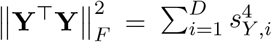. Thus, the objective function (S1) becomes:

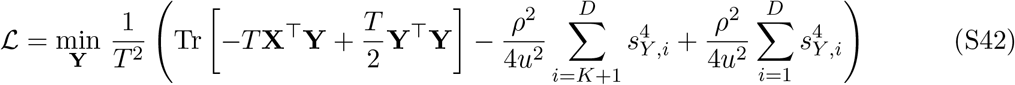

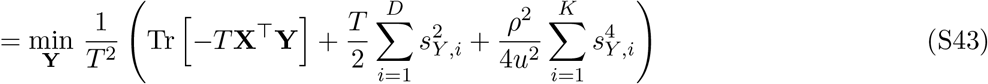

Thus there is a fourth order penalty on the first *K* singular values of **Y**.

We now replace the remaining **X** and **Y** by their SVD:

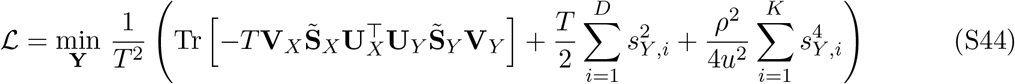

Based on von Neumann trace inequality, given a fixed 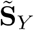, the trace term is minimized when **U***_Y_* = **U***_X_* and **V***_Y_* = **V***_X_*. We are thus left with:

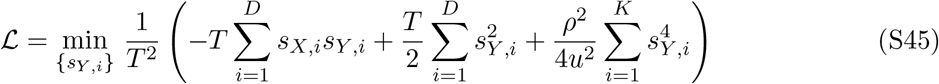

Each *s_Y,i_* can be optimized independently. We take the derivative of (S45) with respect to *s_Y,i_* and equate it to 0. For *i* > *K*, we have *s_Y,i_* = *s_X,i_*. For *i* ≤ *K*

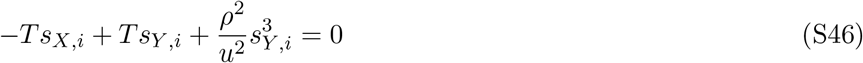

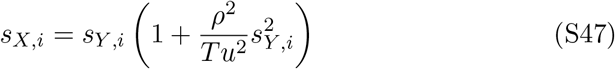

This end the derivation.

#### 4.2 Approach 2

We first find the stationary point of the objective function (S2) in **Y** by taking the partial derivative of 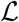 with respect to **Y**:

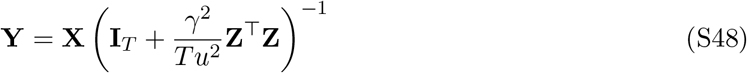

where **I***_T_* is the identity matrix of dimension *T* and replace this solution for **Y** into the objective function 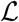:

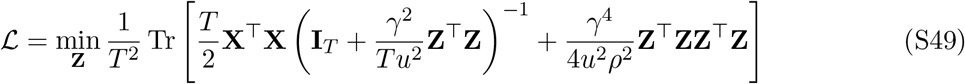

Next we replace **X** and **Z** by their SVD, use the property of the trace Tr(**AB**) = Tr(**BA**) and the property of orthogonal matrices **UU**^⊤^ = **U**^⊤^**U** = **I**:

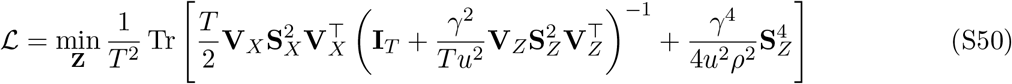

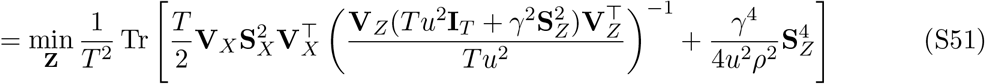

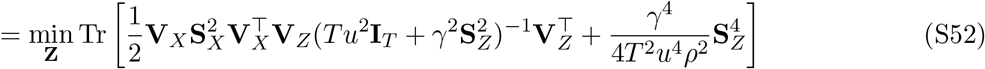

Since **U***_Z_* does not appear in the minimization, it is a free parameter, i.e., it can be any orthogonal matrix. For fixed **S***_Z_*, only the first term in the trace needs to be minimized. One can show that the optimal **V***_Z_* is **V***_Z_* = **V***_X_*: based on von Neumann trace inequality, we know that 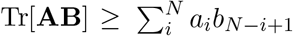 where *a_i_* and *b_i_* are the ordered singular values of **A** and **B**, respectively. Thus, choosing **V***_Z_* = **V***_X_* will give us the lower bound of that inequality. Indeed:

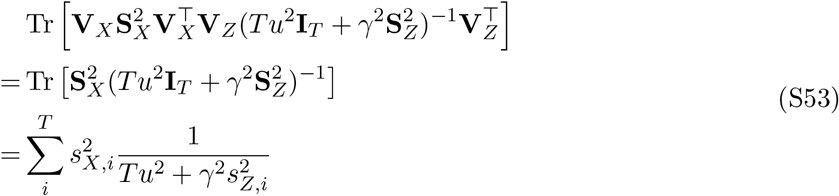

Where *s_X,i_* and *s_Z,i_* are the values on the diagonal of **S***_X_* and **S***_Z_*, respectively. Thus, the highest singular values of 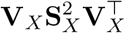 match the lowest singular values of 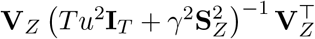, giving us the lower bound of the von Neumann inequality. Equation (S52) can now be simplified to:

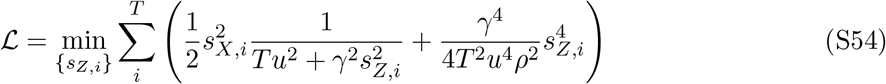

Each *s_Z,i_* can be minimized independently. By construction of SVD, we already have that *s_Z,i_* = 0 for *i* > *K*. We thus consider 1 ≤ *i* ≤ *K*. To simplify notation, we drop the index *i*. We take the derivative of (S54) with respect to *s_Z,i_* and equate it to 0:

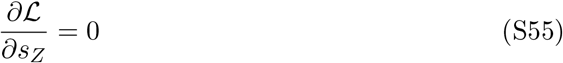

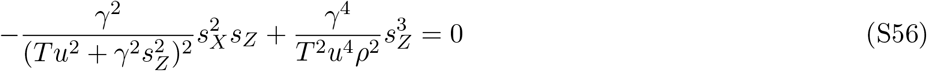

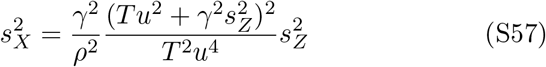

Which leads to, considering that singular values are positive:

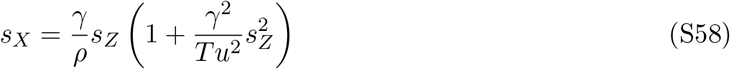

We can now use the obtained solution for **Z** to find the solution for **Y**. We replace **X** and **Z** by their SVD in relation (S48) and use that **V***_X_* = **V***_Z_*:

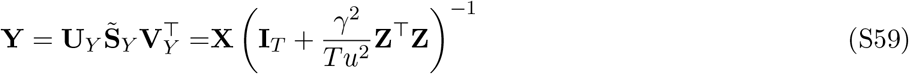

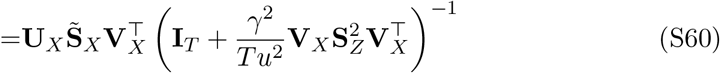

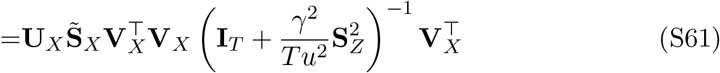

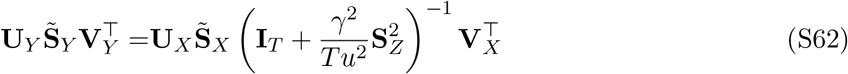

Equating the SVD terms on the left and right sides we obtain **U***_Y_* = **U***_X_* and **V***_X_* = **V***_Y_* and

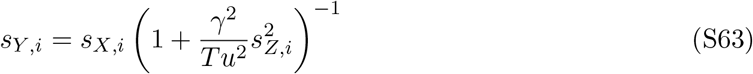

Thus, for *i* > *K*, we have *s_Y,i_* = *s_X,i_* (since *s_Z,i_* = 0), whereas for *i* ≤ *K*: 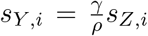 (using relation (S58) to replace *s_X_*). The relation analogous to (S58) is:

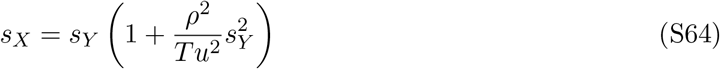

This ends the derivation.

#### 4.3 Effect of *ρ* and relation to SNMF

Having the expression for the output **Y**, we can now describe the effect of *ρ* on the computation. For *ρ* → 0, **Z** → 0, leading to **X** = **Y**, which means that the output is equal to the input and no inhibition is taking place. On the other hand, for *ρ* → ∞, the lowest *D* – *K* singular values of **Y** remain the same, whereas top *K* drop to 0, i.e., the top *K* singular values are totally suppressed.

To better understand the behavior of the circuit for small *ρ* we do a first order expansion in *ρ* of **Y** around **X**, i.e., **Y** = **X** + *ρ***Ξ**. Replacing this expression for **Y** in the objective function (S2), and keeping only the leading terms in *ρ*, the objective function becomes:

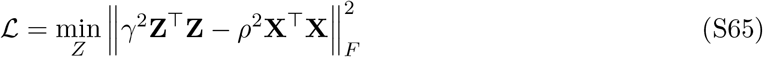

Which corresponds to the basic similarity matching objective function (Pehlevan et al., 2018).

For the non-negative objective function, for small *ρ* we get **Y** = [**X**]_+_ and the objective function simplifies to

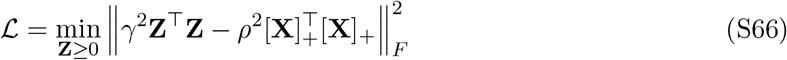

Which corresponds to the symmetric non-negative matrix factorization (SNMF) objective function and can also be implemented online by a neural circuit (Pehlevan & Chklovskii, 2015).

### 5 Relationship between W and M

Here we prove the relationship *ρ*^2^/*γ*^2^**WW**^⊤^ = **M**^2^ for the LC.

One way to obtain this relationship is to start from the circuit dynamics (equations (S17)). The steady state for 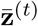 is:

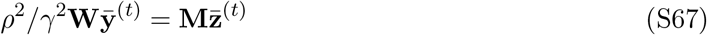

Multiplying by 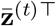 on both sides, taking the average over all samples *t*, and using the definition of **W** and **M** (equation (S13)):

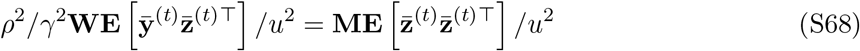

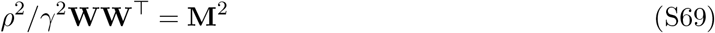

An alternative approach to find the above relationship is to use the definition of **W** and **M** (equation (S13)) and the SVD decomposition of **X, Y**, and **Z**. We write out **W** and **M**:

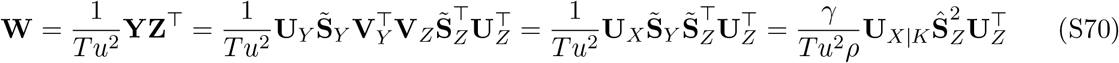

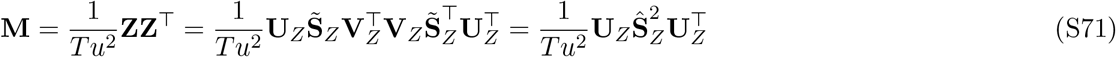

Where we used that **V***_X_* = **V***_Y_* = **V***_Z_* and **U***_X_* = **U***_Y_* are orthogonal matrices and that 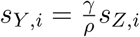 for *i* ≤ *K* and *s_Z,i_* = 0 for *i* > *K*. We call 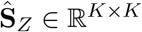 the small square submatrix of the rectangular matrix 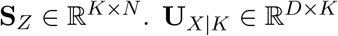 is the submatrix with the first *K* columns of **U***_X_*. Thus:

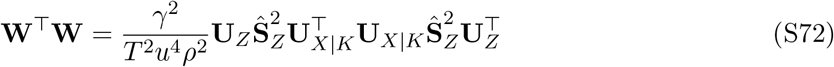

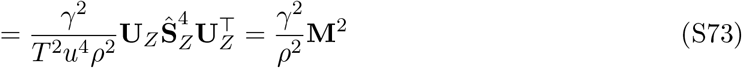

Taking the square root on both sides gives the relationship (2) in the results section.

### 6 Relationship between the statistics of ORN activity and ORN-LN connectivity

Based on the expressions for **W** and **M** (equations (S70) and (S71)) we can write **W** as:

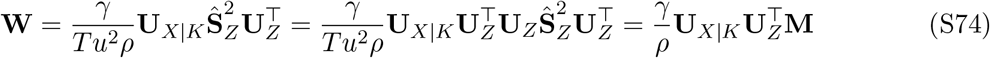

Where we used that 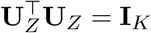. Where 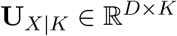 is the submatrix with the first *K* columns of **U***_X_*. As stated above, **U***_Z_* is a free parameter and could be any orthogonal matrix.

In the case of a single LN, **W** is a column vector and corresponds to the first left eigenvector of **X**. For multiple LNs, the column vectors of **W** span the same subspace as the top *K* loading vectors of **X, U***_X|K_*. However, because of the multiplication on the right by 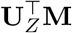, the connections vectors do not necessarily correspond to specific PCA directions and are not orthogonal, but only span the top K-dimensional PCA subspace. Thus, this relation above gives us the relationship between the left eigenvectors of **X, W**, and **M**.

### 7 Decrease of the spread of the spectrum of singular values

Here we show that the coefficient of variation (CV, i.e., the spread) of singular values is smaller at the ORN output (axons) than at the input (somas) in the LC model with the number of ORNs equal to the number of LN. In that case, we have 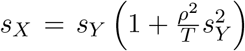. As we have shown, for small *s_X_*, we have *s_Y_* ≈ *s_X_* and for large *s_X_*, we have *s_Y_* ≈ (*T/ρ*^2^*s_X_*)^1/3^. We call *X* a positive random variable. We will show that for a 0 < *α* < 1, *CV*(*X*) ≥ *CV*(*X^α^*), which mimics the case we have.

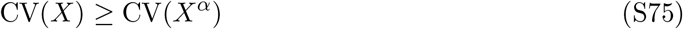

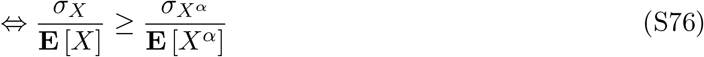

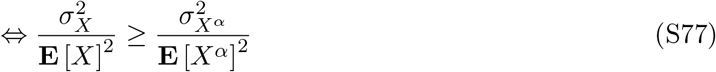

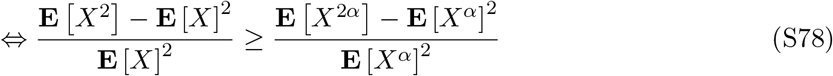

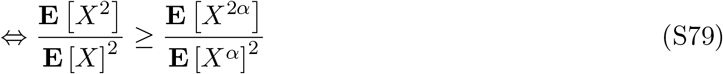

The last inequality can be proven by using Hölder’s inequality twice. First:

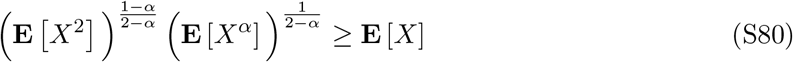

which leads to:

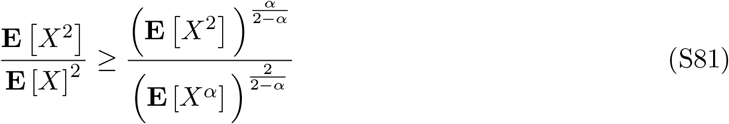

and second:

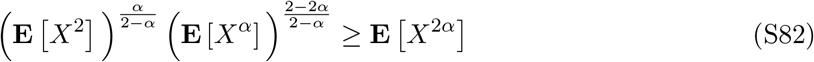

which leads to:

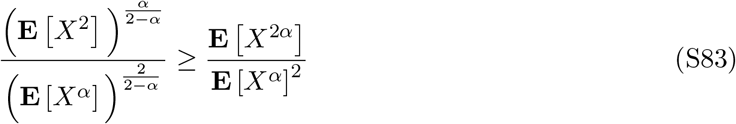

Combining inequalities (S81) and (S83) proves inequality (S79) and ends the proof.

